# Identification of Polycomb Repressive Complex 1 and 2 Core Components in Hexaploid Bread Wheat

**DOI:** 10.1101/703546

**Authors:** Beáta Strejčková, Radim Čegan, Ales Pecinka, Zbyněk Milec, Jan Šafář

**Affiliations:** The Czech Academy of Sciences, Institute of Experimental Botany (IEB), Centre of the Region Haná for Biotechnological and Agricultural Research (CRH), 78371 Olomouc, Czech Republic; Department of Plant Developmental Genetics, Institute of Biophysics, Academy of Sciences of the Czech Republic, 61200 Brno, Czech Republic

## Abstract

Polycomb repressive complex 1 and 2 play important roles in epigenetic gene regulation by posttranslationally modifying specific histone residues. Polycomb repressive complex 2 is responsible for the trimethylation of lysine 27 on histone H3, while Polycomb repressive complex 1 catalyzes the monoubiquitination of histone H2A at lysine 119. Although these biochemical functions are evolutionarily conserved, studies in animals and plants, mainly *Arabidopsis thaliana*, showed that specific subunits have evolved into small gene families, with individual members acting at different developmental stages or responding to specific environmental stimuli. However, the evolution of polycomb group gene families in monocots, particularly those with complex allopolyploid origins, is unknown. Here, we present the *in silico* identification of the Polycomb repressive complex 1 and 2 subunits in allohexaploid bread wheat, the reconstruction of their evolutionary history and a transcriptional analysis over a series of 33 developmental stages. The identification and chromosomal location of the Polycomb repressive complex 1 and 2 core components in bread wheat may enable a deeper understanding of developmental processes, including vernalization in commonly grown winter wheat.

## INTRODUCTION

The regulation of gene expression in higher organisms includes a wide range of mechanisms acting at the transcriptional, posttranscriptional and posttranslational levels. More complex regulation that is required to coordinate proper gene activity also includes regulation by chromatin remodeling via histone modifications (methylation, acetylation, phosphorylation and ubiquitination), which lead to specific chromatin changes. Prominent posttranslational changes are histone modifications, which occur on particular amino acid residues. Methylation of lysine 4 on histone H3 (H3K4me) is mainly associated with transcriptional activation, while the di- and trimethylation of lysines 9 and 27 (H3K9me2 and H3K27me3, respectively) leads to transcriptional repression (Wu *et al*. 2009). H3K9me2, together with small double stranded RNAs and DNA hypermethylation, contributes to the silencing of repetitive DNA sequences (Matzke and Mosher, 2014; Fultz et al., 2015). The repressive epigenetic regulatory processes of genes are usually controlled by Polycomb group proteins (PcG), which are, at the basic level, evolutionarily conserved among plants and animals (Mozgova and Hennig 2015). Polycomb repressive complex 1 (PRC1) and 2 (PRC2) are two of the main complexes involved in developmental gene regulation and were initially identified in *Drosophila melanogaster* (reviewed in (Margueron and Reinberg 2011; Schwartz and Pirrotta 2013; Mozgova and Hennig 2015). Traditionally, PRC1 and PRC2 have been suggested to work in a hierarchical PRC2 → PRC1 manner (Wang *et al*. 2004), but recently, a PRC2-independent function of PRC1 has been suggested (Kahn *et al*. 2016; Dorafshan *et al*. 2017) . According to the hierarchical model, PRC2 binds to specific DNA sequence motifs called Polycomb response elements (PRE) and trimethylates H3 at lysine 27 (H3K27me3) in the nearby nucleosomes. This recruits PRC1, which catalyzes the monoubiquitination of histone H2A (H2AK119u1) and stabilizes H3K27me3 modification by chromatin remodeling (Endoh *et al*. 2012). The PRC2:PRC1-independent model proposes that PRC1 and PRC2 have their own specific adaptor proteins that bind the PRE, and consequently, PRC1/2 are independently recruited via interactions with the particular adaptor proteins (Dorafshan *et al*. 2017).

*Drosophila* PRC1 contains four core components, Polycomb (Pc), Polyhomeotic (Ph), Posterior sex combs (Psc) and Sex combs extra (Sce), although a fifth component, Sex combs on midleg (Scm), has also been reported (reviewed in (Schwartz and Pirrotta 2013)). The presence of PRC1 has been unclear in plants until RING-finger proteins were described in *Arabidopsis* (Xu and Shen 2008; Chen *et al*. 2010). In *A. thaliana*, LIKE HETEROCHROMATIN PROTEIN1 (AtLHP1) substitutes the role of Pc (Turck *et al*. 2007). With its chromo domain, LHP1 recognizes and binds histone H3 methylated lysine 27 (H3K27me3) (Zhang *et al*. 2007). *A. thaliana B LYMPHOMA Mo-MLV INSERTION REGION 1 HOMOLOG* (*AtBMI1A to C*) are three homologs of Psc, and *REALLY INTERESTING NEW GENE1* (*AtRING1A* and *AtRING1B*) are two homologs of Sce (reviewed in (Chen *et al*. 2016)). No Ph homolog has been identified in plants to date (Bemer and Grossniklaus 2012). However, plant-specific proteins related to PRC1, such as *A. thaliana* EMBRYONIC FLOWER1 (AtEMF1) (Calonje *et al*. 2008) or *A. thaliana* VERNALIZATION1 (AtVRN1) (Mylne *et al*. 2006) were suggested. EMF1 is involved in the control of shoot architecture and flowering in *Arabidopsis* (Aubert *et al*. 2001) and interacts with the AtBMI1 and AtRING1 homologs of PRC1 (Bratzel *et al*. 2010, 2012). In contrast, there is no report on the interactions between AtVRN1, which is involved in vernalization in *Arabidopsis* (Levy *et al*. 2002), and other PRC1 components (Berke and Snel 2015); thus, there is no consensus regarding whether VRN1 is a component of PRC1. Recently, an alternative complex with a PRC1-like function was reported (Li *et al*. 2018). In *Arabidopsis,* two homologous BAH (Bromo-adjacent homology) domain–containing proteins form a plant-specific complex with EMBRYONIC FLOWER1 (AtEMF1), and this BAH–EMF1 complex reads and effects the H3K27me3 mark and mediates genome-wide transcriptional repression. A homolog of a BAH-domain protein was also found in monocots (rice), which may indicate its conservation in flowering plants (Li *et al*. 2018). Genes encoding PRC1 subunits have also been reported in monocots, e.g., *Zea mays* and *Oryza sativa* (Berke and Snel 2015), but not in agronomically important temperate zone cereals, such as wheat or barley.

The PRC2 complex is formed by four subunits: Enhancer of zeste [E(z)], Extra sex combs (Esc), Suppressor of zeste 12 [Su(z)12] and WD protein p55 (Bantignies and Cavalli 2011); however, similarly to PRC1, an additional fifth core component (Jing) has been suggested in *Drosophila* (Schwartz and Pirrotta 2013). In plants, PRC2 has been thoroughly studied in *Arabidopsis* (reviewed in (Mozgova and Hennig 2015). The catalytic activity of PRC2 is histone methylation associated with the SET domain in E(z). Three E(z) homologs have been described: CURLY LEAF (CLF) (Chanvivattana 2004), SWINGER (SWN) (Goodrich *et al*. 1997) and MEDEA (MEA) (Grossniklaus *et al*. 1998). Similarly, three homologs of Su(z) have been identified: REDUCED VERNALIZATION RESPONSE2 (VRN2) (Gendall *et al*. 2001), EMBRYONIC FLOWER2 (EMF2) (Yoshida 2001) and FERTILIZATION INDEPENDENT SEED2 (FIS2) (Luo *et al*. 1999a). The *ESC* homolog *FERTILIZATION INDEPENDENT ENDOSPERM* (*FIE*) is present as a single gene, while the WD40 p55 homolog (*MULTICOPY SUPPRESSOR OF IRA1*, *MSI*) was found to have five genes (*MSI1 to MSI5*) in *Arabidopsis* (Hennig 2003). Each of the *Arabidopsis E(z)* and *Su(z)* homologs functions at different developmental stages (reviewed in (Derkacheva and Hennig 2014). The E(z) homolog MEA is active during early endosperm development (Köhler *et al*. 2003), while SWN and CLF play a role in vegetative development and vernalization. The initiation of flowering after vernalization is controlled by the flowering repressor *FLOWERING LOCUS C* (*FLC*) (Michaels and Amasino 1999; Sheldon *et al*. 1999). It was shown that during vernalization, the H3K27me3 level increases and gradually silences *FLC* (Angel *et al*. 2011), and *FLC* is switched off completely at the end of the cold period (Sheldon *et al*. 2000). This status is reset in the next generation, and thus, plants must undergo vernalization to flower.

In *Arabidopsis*, the *clf swn* double mutant completely loses H3K27me3, which indicates the possible inactivation of PRC2 (Lafos *et al*. 2011). However, *clf swn* plants form only callus-like structures with occasional somatic embryos (He *et al*. 2012). The Su(z) homolog FIS takes part in the regulation of the female gametophyte and seed development (Chaudhury *et al*. 1997), but the Su(z) homolog EMF2 controls the transition to flowering (Yang *et al*. 1995). Grass PRC2 homologs have been identified *in silico* in maize, rice and barley (Phillips *et al*., 2002; Thakur *et al*., 2003; Hennig *et al*., 2005; Haun *et al*., 2007; Chen *et al*., 2009; Luo *et al*., 2009; Kapazoglou *et al*., 2010). Their function has been associated mainly with seed and endosperm development (Kapazoglou *et al*. 2010; Tonosaki and Kinoshita 2015); for a detailed summary, see (Butenko and Ohad, 2011). Kapazoglou *et al*. (2010) identified the barley PRC2 homologs *HvFIE*, *HvE(z)*, *HvSu(z)12a* and *HvSu(z)12b*, but *p55* has not been found.

Recently, (Lomax *et al*. 2018) identified a *Brachypodium distachyon* mutant without vernalization requirements. A mutation in *Enhancer of zeste-like* (*EZL1*), an ortholog of *A. thaliana CLF*, caused an overall reduction in H3K27me3 and H3K27me2 at *B. distachyon VERNALIZATION1* (*BdVRN1*) and, consequently, earlier flowering without vernalization. A significant reduction in H3K27me3 levels in several regions of *TaVRN1* during vernalization was also reported in the bread wheat *Triticum aestivum*, which was positively correlated with the length of the cold period (Xiao *et al*. 2014). This could indicate an important role of PRC2-mediated H3K27me3 deposition in the process of vernalization in grasses.

Despite the socioeconomic importance of bread wheat, the reference genome sequence has been published only for the model cultivar Chinese Spring (The International Wheat Genome Sequencing Consortium (IWGSC) *et al*. 2018). Bread wheat (2n = 6x = 42) is a recently formed allohexaploid with a vast nuclear genome (16 974 Mb/1C, (Bennett and Smith 1991) assembled from three homoeologous subgenomes (A, B and D). Thus, the deep analyses of genes and their biochemical pathways are lacking behind those of other crops and model plant species, such as *A. thaliana*.

Here, we report the identification and chromosomal location of bread wheat genes coding the individual subunits of PRC2 and PRC1. We analyzed the mRNA levels of individual genes at different developmental stages and showed sequence conservation among other *Triticeae* species, such as *Triticum urartu*, *Aegilops tauschii* and *Triticum dicoccoides*, using a phylogenetic approach. We also discuss the putative role of PRC2 and PRC1 in the vernalization process in bread wheat.

## MATERIALS AND METHODS

### *In silico* PcG component identification

*T. aestivum* PcG component protein sequences were obtained by BLAST searches of the *T. aestivum* genome in Ensembl Plants (http://plants.ensembl.org/index.html) using *A. thaliana* protein sequences. Not all wheat full protein sequences were available in the database. Therefore, some sequences were *in silico* reconstructed from the genomic sequences according to the *T. aestivum* reference (cultivar Chinese Spring) obtained from Ensembl Plants by local blastn with genomic data of *T. urartu* and *Ae. tauschii*. Data for *T. dicoccoides* were obtained from Ensembl Plants. The obtained nucleotide sequences were aligned to the *T. aestivum* sequence by MAFFT multiple aligner (version 1.3.3) in Geneious 8.1.9 software https://www.geneious.com. After alignment of genomic sequences, coding sequence (CDS) regions were extracted and translated into proteins. Some genomic sequences were not well assembled, and thus, a sequence corresponding to the reference was sometimes scattered to several scaffolds/contigs. Such genes were reconstructed by extracting partial sequences from several scaffolds, concatenating the CDS regions and translating them into proteins.

Protein sequences for *Hordeum vulgare* were obtained from GenBank https://www.ncbi.nlm.nih.gov/ and barley DB (Monat *et al*. 2019) ; proteins for *B. distachyon*,

*Helianthus annuus*, *Nicotiana attenuata*, *Oryza sativa japonica*, *Oryza sativa indica, Populus trichocarpa*, *Solanum lycopersicum* and *Z. mays* were obtained from UniProt (https://www.uniprot.org/) and Ensembl Plants. All sequences used in the phylogenetic studies are provided in Supplementary Table 3.

Reciprocal BLAST searches of identified wheat PcG proteins were performed against the *A. thaliana* database TAIR10 within EnsemblPlants (https://plants.ensembl.org/Arabidopsis_thaliana/Info/Index) to validate the results.

### Phylogenetic analysis

Protein alignments for phylogenetic analysis were conducted in MEGA X (Kumar *et al*. 2018) by ClustalW. For all genes in PRC1 and PRC2 complexes, the evolutionary history was inferred by using the maximum likelihood method and JTT matrix-based model (Jones *et al*. 1992) in MEGA X (Kumar *et al*. 2018) The bootstrap consensus tree inferred from 1000 replicates (Felsenstein 1985) is taken to represent the evolutionary history of the taxa analyzed (Felsenstein 1985). Sequences of *Drosophila* PcG proteins were used as outgroups for all trees besides EMF1 where *Arabidopsis* sequence was used as outgroup. All phylogenetic trees were rooted in the outgroup except E(z), which were rooted at the midpoint.

### Transcriptomic analysis

The RNA-seq database “expVIP” http://www.wheat-expression.com was used as a data source for the expression analysis of individual PcG core subunits (Borrill *et al*. 2016; Ramírez-González *et al*. 2018). We used data collected from roots, leaves/shoots, spikes and grains of spring wheat cultivar Azhurnaya at 58 different time points, corresponding to a total of 22 tissues or organs (Supplementary Table 2). Heatmaps were constructed in R software (https://www.r-project.org/) using gplots, heatmap3 and RColorBrewer packages. Both the genes and the developmental stages were clustered based on the similarity of their mRNA amounts at different experimental points.

### Protein domain identification

The SMART http://smart.embl.de/ (in mode normal SMART) (Letunic and Bork 2018) and PFAM http://pfam.xfam.org/ (El-Gebali *et al*. 2019) protein databases were used to predict conserved protein domains of the PRC2 and PRC1 components of *A. thaliana*, *H. vulgare*, *T. dicoccoides* and *T. aestivum*. A multiple sequence alignment of all found homologous proteins for each PRC2 and PRC1 subunit of *A. thaliana*, *H. vulgare*, *T. dicoccoides* and *T. aestivum* was carried out by using MAFFT *v7.388* (Katoh et al. 2002; Katoh and Standley 2013).

## RESULTS

### *In silico* identification of wheat PRC2 and PRC1 core components

Using protein sequences of the *Arabidopsis* PcG homologs, we identified all wheat components with their respective chromosomal locations. As expected, homoeologs of individual components on all three wheat subgenomes A, B and D were also located. The bread wheat components have been designated with the prefix “Ta” representing *Triticum aestivum* followed by A, B or D to determine the subgenome location. If additional entries were identified on a different chromosome or the same chromosome but at different positions, the respective number was added to distinguish between individual paralogs, for example, *TaSu(z)-2A1* (Table 1 and Figure 1A). The chromosomal positions were validated using the available reference genomes of *T. urartu* (2n = 2x = 14)*, T. dicoccoides* (wild emmer wheat, 2n = 4x = 28, accession Zavitan) and *H. vulgare* (2n = 2x = 14, cultivar Morex) (Supplementary Table 1).

**Table 1.**
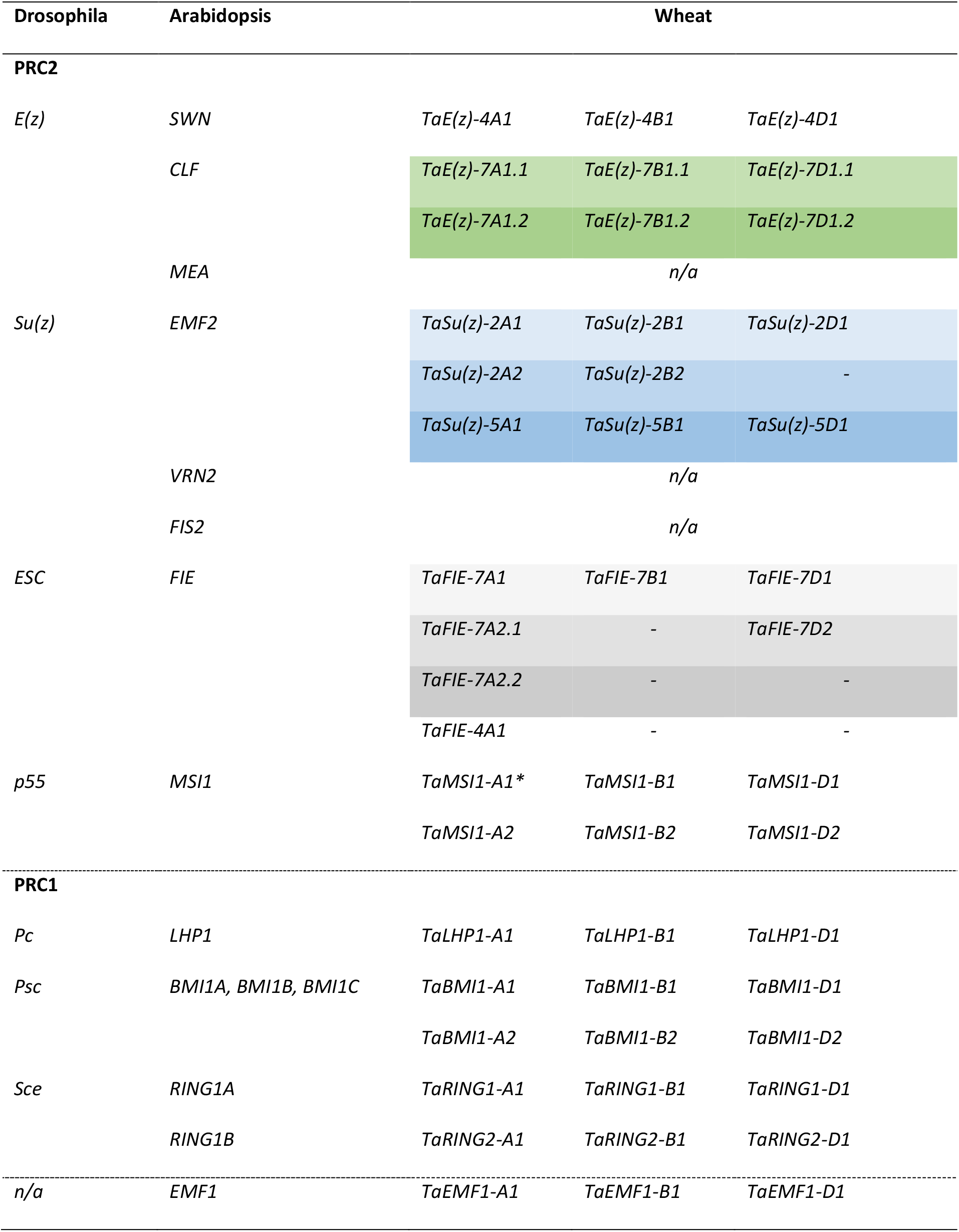
Polycomb group core components. The table shows genes of PRC2 and PRC1 previously reported in *Drosophila*, *Arabidopsis* and those identified in bread wheat. Each column in wheat contains A, B, and D subgenome homoeologs. Shades of the same color represent putative paralogs. EMF1 is a plant-specific PRC1-related component that is not present (n/a) in Drosophila. A dash (-) indicates that no homolog was identified. * indicates the gene not assigned to any chromosome based on a BLAST search. The accession numbers of the respective wheat PcG components are listed in Supplementary Table 1.

| Drosophila | Arabidopsis | Wheat |  |  |
| --- | --- | --- | --- | --- |
| PRC2 |  |  |  |  |
| E(z) | SWN | TaE(z)-4A1 | TaE(z)-4B1 | TaE(z)-4D1 |
|  | CLF | TaE(z)-7A1.1 | TaE(z)-7B1.1 | TaE(z)-7D1.1 |
|  |  | TaE(z)-7A1.2 | TaE(z)-7B1.2 | TaE(z)-7D1.2 |
|  | MEA | n/a |  |  |
| Su(z) | EMF2 | TaSu(z)-2A1 | TaSu(z)-2B1 | TaSu(z)-2D1 |
|  |  | TaSu(z)-2A2 | TaSu(z)-2B2 | - |
|  |  | TaSu(z)-5A1 | TaSu(z)-5B1 | TaSu(z)-5D1 |
|  | VRN2 | n/a |  |  |
|  | FIS2 | n/a |  |  |
| ESC | FIE | TaFIE-7A1 | TaFIE-7B1 | TaFIE-7D1 |
|  |  | TaFIE-7A2.1 | - | TaFIE-7D2 |
|  |  | TaFIE-7A2.2 | - | - |
|  |  | TaFIE-4A1 | - | - |
| p55 | MSI1 | TaMSI1-A1* | TaMSI1-B1 | TaMSI1-D1 |
|  |  | TaMSI1-A2 | TaMSI1-B2 | TaMSI1-D2 |
| PRC1 |  |  |  |  |
| Pc | LHP1 | TaLHP1-A1 | TaLHP1-B1 | TaLHP1-D1 |
| Psc | BMI1A, BMI1B, BMI1C | TaBMI1-A1 | TaBMI1-B1 | TaBMI1-D1 |
|  |  | TaBMI1-A2 | TaBMI1-B2 | TaBMI1-D2 |
| Sce | RING1A | TaRING1-A1 | TaRING1-B1 | TaRING1-D1 |
|  | RING1B | TaRING2-A1 | TaRING2-B1 | TaRING2-D1 |
| n/a | EMF1 | TaEMF1-A1 | TaEMF1-B1 | TaEMF1-D1 |

**Figure 1.**
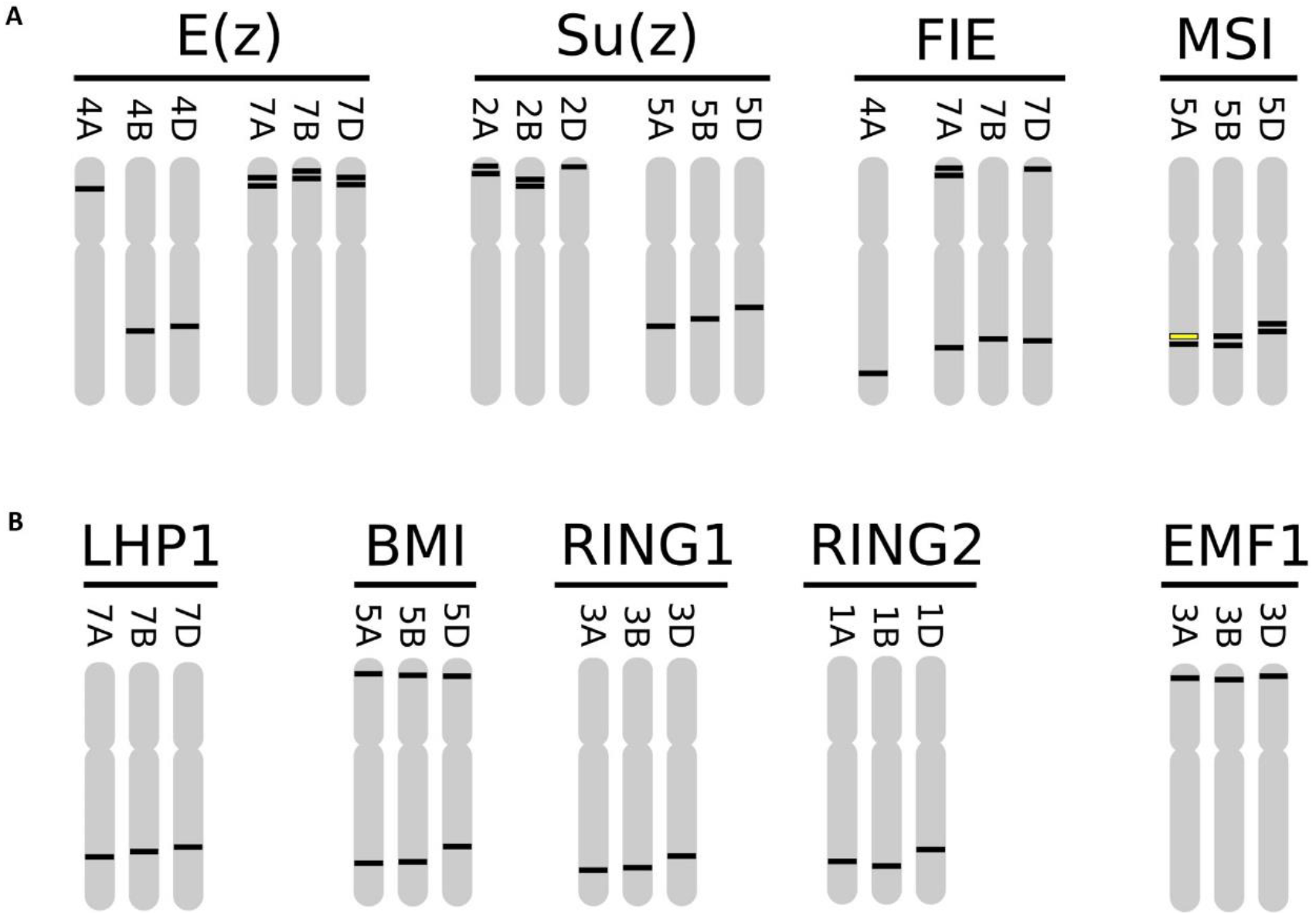
Schematic representation of PRC2 (A) and PRC1 (B) component positions on the chromosomes of bread wheat. The scheme shows the locations of individual wheat homologs based on a BLAST search. The yellow line on chromosome 5A for the MSI homolog shows the gene TraesCSU02G072700, which was not anchored to any chromosome originally. The depicted positions are only illustrative and not in scale due to the small size of the picture.

*Enhancer of zeste* [*E(z)*] is located on chromosomes 4 and 7 (Table 1 and Figure 1A). On chromosome 4, *E(z)* was found on the short arm [*TaE(z)-4A1*] and on the long arm [*TaE(z)- 4B1*, *TaE(z)-4D1*], while on chromosome 7, *E(z)* was found on the short arm (Supplementary Table 1). The position of *TaE(z)-4A1* on the short arm of chromosome 4A corresponds with the pericentric inversion reported in hexaploid wheat(Hernandez *et al*. 2012; The International Wheat Genome Sequencing Consortium (IWGSC) *et al*. 2018). On chromosome 7, two paralogs on the respective short arm were found, separated by only a tens of kb region, suggesting that they originated from a local gene duplication event (Supplementary Table 1). Furthermore, these paralogs differed by approximately ≈ 85 amino acids as a result of multiple insertions and deletions (indels), with the longest indel 137 amino acids in length (Supplementary Figure S1D).

Kapazoglou *et al*. (2010) reported several *Suppressor of zeste* [Su(z)] homologs in barley, located on chromosomes 2H and 5H. Similarly, we found wheat homologs on chromosomes 2 and 5. Interestingly, there were two homologs identified on chromosomes 2AS and 2BS, but only one on 2DS (Table 1 and Figure 1A). All three homoeologs of group 5 were located on the long arm. Bread wheat diploid progenitor *T. urartu* has only an A genome, and we identified two homologs on the short arm of chromosome 2 at positions ≈ 1.5 Mb and ≈ 2.4 Mb and another one on the long arm of chromosome 5. Wild emmer wheat accession Zavitan also carries two homologs on 2AS and one on 2BS together with homologs on 5AL and 5BL (Supplementary Table 1).

Two proteins coded by genes *TaSu(z)-2A2* and *TaSu(z)-2B2* carry an insertion of ≈ 30 amino acids. This insertion was also found in proteins encoded by the *TRIDC2AG000370.14* gene in *T. dicoccoides* and by the *H. vulgare* gene *HORVU.MOREX.r2.2HG0078790.1* located on chromosome 2 (Supplementary Figure S1G).

The Esc subunit reported in *Drosophila* was designated *FERTILIZATION INDEPENDENT ENDOSPERM1* (*HvFIE1*) in barley (Kapazoglou *et al*. 2010), so we followed this style and designated the wheat homologs *TaFIE*. We found two homologs on 7AS (*TaFIE-7A2.1* and *TaFIE-7A2.2*) and one on 7AL (*TaFIE-7A1*) chromosome (Table 1, Figure 1A, and Supplementary Table 1A). Chromosome 7D has one gene located on the short arm (*TaFIE-7D1*) and one gene on the long arm (*TaFIE-7D2*). Initially, no 7B homolog was located using the reference sequence of Chinese Spring by IWGSC. Surprisingly, a paralog was located in the distal part of the long arm of chromosome 4. This corresponds with the fact that this region of chromosome 4 contains a part of chromosome 7B (Hernandez *et al*. 2012). Reciprocal BLAST with the 4AL homolog (TaFIE-4A1) showed high similarity with genes previously located on 7AL/7BL in Zavitan and with the barley gene on the 7H chromosome. The predicted barley protein was annotated as FIE (Mascher *et al*. 2017; Monat *et al*. 2019). Later, we identified the 7BL homolog TRIAE_CS42_7BL_TGACv1_580129_AA1912160.1 using a BLAST search in the Ensembl plant database using data from wheat genome assembly by TGAC (Clavijo *et al*. 2017) (Supplementary Table 1).

The p55 subunit carries WD40 domains (same as FIE) and was designated MSI1 (MULTICOPY SUPPRESSOR OF IRA1) in *Arabidopsis*. In bread wheat, two orthologs (*TaMSI1)* were located on each chromosome of group 5 with one exception: one of the best BLAST results was not anchored to any chromosome (TraesCSU02G072700). A comparison with the sequences of *T. urartu* and *T. turgidum* revealed high identity with the 5AL chromosome; therefore, we designated this unassigned accession *TaMSI1-A1,* suggesting its location to chromosome 5A (Table 1 and Supplementary Table 1).

The location of wheat PRC1 components was more complicated, as they have not been described in cereals thus far, which made validation of the results more difficult. Therefore, we used the reference sequence of *H. vulgare* containing an annotation of predicted proteins.

*LIKE HETEROCHROMATIN PROTEIN1* (*LHP1*) wheat homoeologs were located on the long arm of chromosome 7. *BMI1* homologs were located on both short and long arms of chromosome 5. *Arabidopsis* has three *BMI1* homologs (*AtBMI1A* to *AtBMI1C*), but BLAST of *AtBMI1A* and *AtBMI1B* identified the same genes in wheat located on the long arm of chromosome 5. Surprisingly, a BLAST search of *AtBMI1c* identified not only the same wheat homologs but also other paralogous genes located on the short arm. The genes on the short arm corresponded to the position of the barley gene, which was also located on the short arm of chromosome 5H. This gene was annotated as Ubiquitin ligase DREB2A-INTERACTING PROTEIN2 (DRIP2, synonym for BMI1) (Monat *et al*. 2019) and corresponded to the *Arabidopsis* designation. The genes on the long arm corresponded with the position of the barley gene annotated also as Ubiquitin ligase DRIP2 (Monat *et al*. 2019) and located on the long arm of chromosome 5H.

The RING1 homologs were found on the long arm of all three chromosomes of group 3, and RING2 was located on the long arm of all three chromosomes of group 1.

The wheat homolog TaEMF1 was not identified when the *Arabidopsis* protein sequence was used in a BLAST search. Homologous proteins with genes located on chromosomes 3A, 3B and 3D were found when the EMF1 protein sequence of *Z. mays* was used (Berke and Snel 2015). The positions of these genes correlated with the location of *HvEMF1* in barley, suggesting that they could be homologs of *AtEMF1*.

We also identified the main protein domains for individual PcG wheat components (Figure 2). The comparison of bread wheat with *Arabidopsis*, *H. vulgare* and *T. dicoccoides* showed high domain conservation, which further supported the accuracy of the wheat homolog identification.

**Figure 2.**
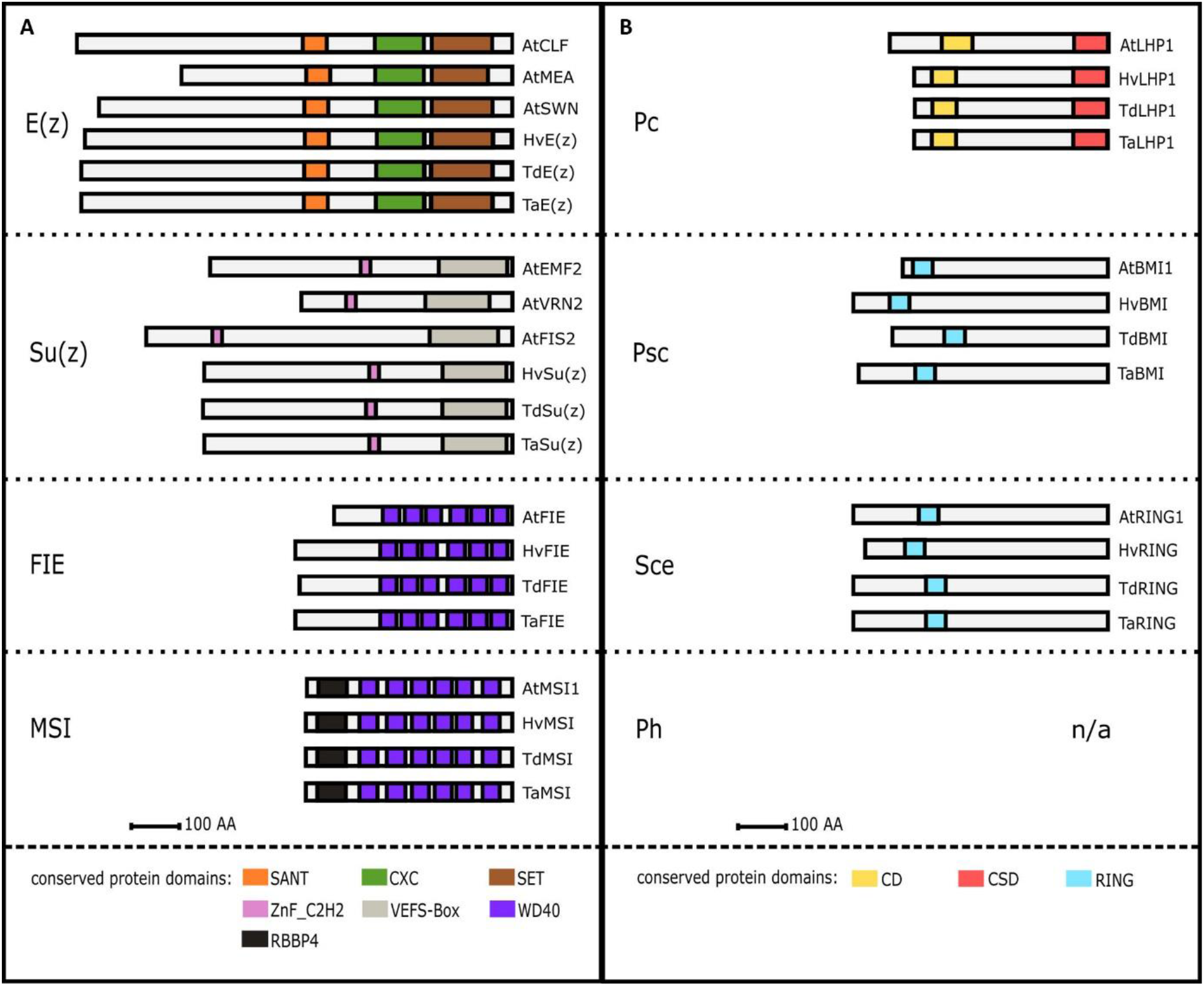
Schematic representation of the conserved protein domain architecture of Polycomb group (PcG) complexes. The *in silico* identification of the PRC2 and PRC1 core components in hexaploid wheat was supported by protein alignment with known homologs from *Arabidopsis* and barley PRC2 and PRC1 and by prediction of main functional protein domains. Homologs of the PRC2 (A) and PRC1 (B) core subunits share highly conserved protein domains among *Arabidopsis thaliana* (At), *Hordeum vulgare* (Hv), *Triticum dicoccoides* (Td) and *Triticum aestivum* (Ta). n/a - not available.

### Phylogenetic analysis

Phylogenetic trees of both PRC2 and PRC1 wheat components have been constructed to reveal the evolutionary relationships among *Arabidopsis*, barley, rice, maize, all bread wheat homologs and bread wheat progenitors. Phylogenetic analysis showed that wheat E(z) homologs, located on chromosomes 4 and 7, fell into separate clades, one including AtSWN and the other AtCLF, respectively. This suggests that *E(z)* genes on wheat chromosome 4 are putative orthologs of *AtSWN*, while genes on chromosome 7 are putative orthologs of *AtCLF* (Figure 3A).

**Figure 3.**
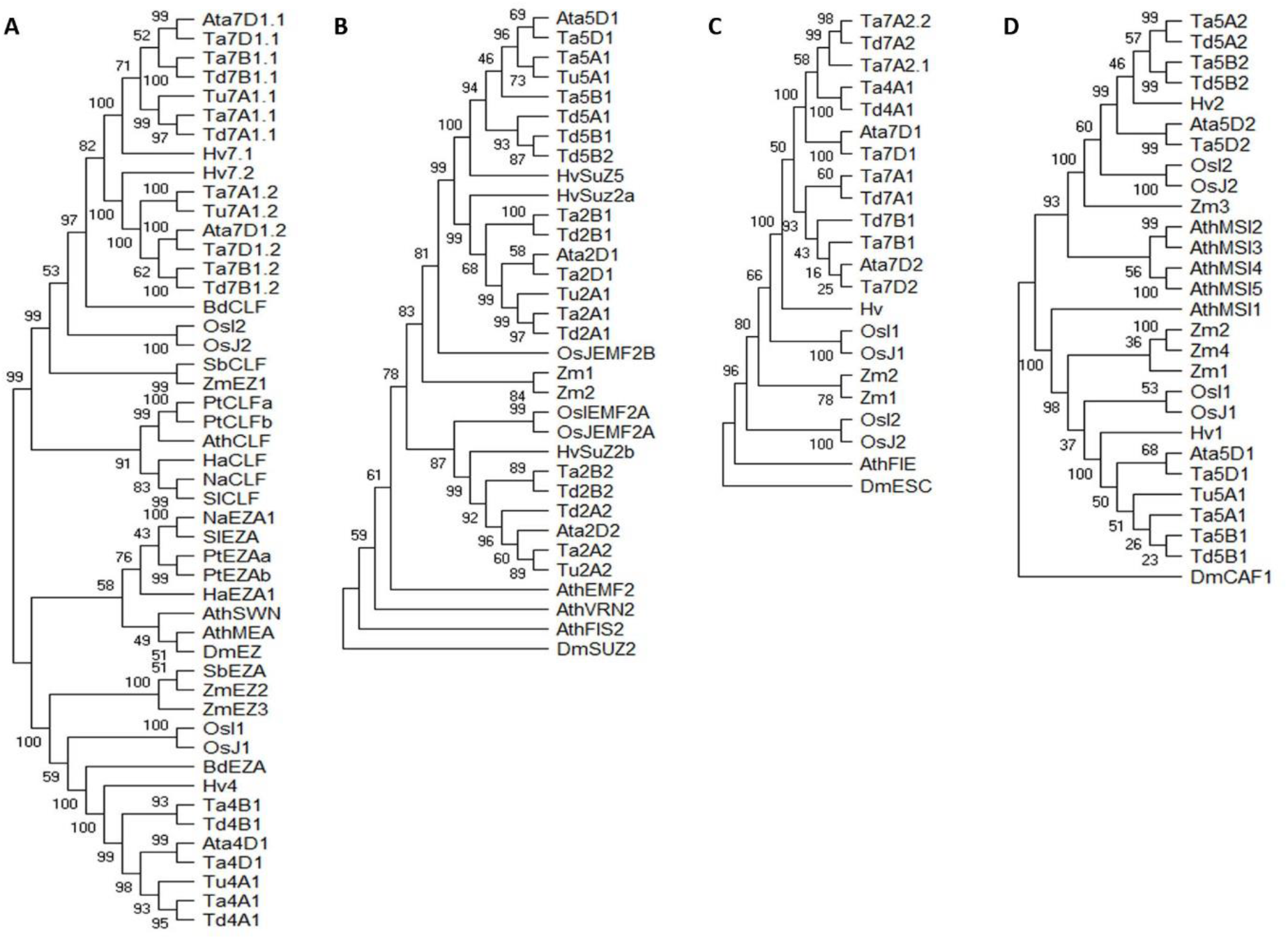
Phylogenetic analysis of the plant PRC2 components E(z) (A), Su(z) (B), FIE (C) and MSI (D). The analysis was performed using the maximum likelihood method and JTT matrix-based model in MEGA X. The bootstrap consensus tree was inferred from 1000 replicates. E(z) tree is midpoint rooted. Su(z), FIE and MSI trees are rooted in the outgroup *Drosophila melanogaster* (Dm). *Aegilops tauschii* (Ata), *Arabidopsis thaliana* (Ath), *Brachypodium distachyon* (Bd), *Helianthus annuus* (Ha), *Nicotiana attenuata* (Na), *Populus trichocarpa* (Pt), *Solanum lycopersicum* (Sl), *Sorghum bicolor* (Sb), *Hordeum vulgare* (Hv), *Oryza sativa indica* (OsI), *Oryza sativa japonica* (OsJ), *Triticum aestivum* (Ta), *Triticum dicoccoides* (Td), *Triticum urartu* (Tu) and *Zea mays* (Zm).

*Su(z)* genes were found on chromosomes 2 and 5. The genes on chromosome 2 clustered in one clade, while genes on chromosome 5 clustered into the second clade. The phylogenetic analysis suggests that all *Su(z)* are orthologous to *AtEMF2* (Figure 3B).

Homologs of *FIE* were located on chromosome 7, but the best BLAST hit was on chromosome 4A. Interestingly, the homolog on the 4AL chromosome (*TaFIE-4A1*) fell into the same clade with the 7AS chromosome homologs (*TaFIE-7A2.1* and *TaFIE-7A2.2*) and not in the clade with the 7AL homolog (Figure 3C).

*MSI* homologs were located in two positions on the long arm of chromosome 5, except for *TraesCSU02G072700*, which was not assigned to any chromosome (Supplementary Table 1). However, phylogenetic clustering of this unanchored gene in the same clade together with *TaMSI1-B1* and *TaMSI1-D1* implies that it may represent the *TaMSI* copy located on the 5A chromosome (Table 1 and Figure 1A).

The phylogenetic analysis of PRC1 components was unremarkable: wheat LHP1 homologs clustered according to subgenomes A, B and D. Although *Arabidopsis* has three BMI homologs, wheat BMI1 homologs fell into only two clades. This was in agreement with our findings based on the alignment (Supplementary Table 1). RING homologs clustered into two clades according to their location on chromosomes 1 and 3 (Figure 4B).

**Figure 4.**
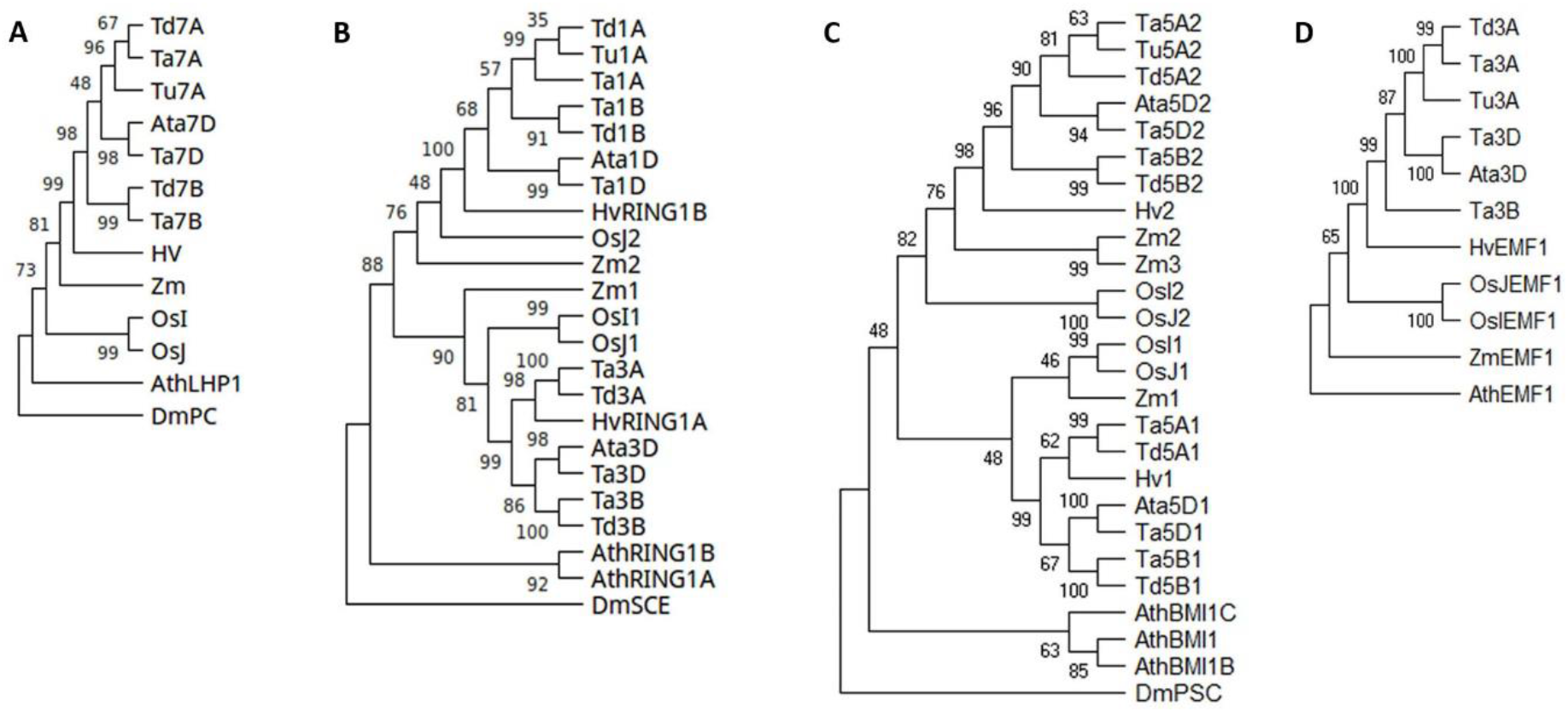
Phylogenetic analysis of the plant PRC1 components LHP1 (A), RING1 (B), BMI1 (C) and EMF1 (D). The analysis was performed using the maximum likelihood method and JTT matrix-based model in MEGA X. The bootstrap consensus tree was inferred from 1000 replicates. Trees are rooted in the outgroup *Drosophila melanogaster* (Dm), with the exception of the EMF1 tree, which is rooted in *Arabidopsis thaliana* (Ath). *Aegilops tauschii* (Ata), *Hordeum vulgare* (Hv), *Oryza sativa indica* (OsI), *Oryza sativa japonica* (OsJ), *Triticum aestivum* (Ta), *Triticum dicoccoides* (Td), *Triticum urartu* (Tu) and *Zea mays* (Zm).

### RNA-seq analysis suggests conserved transcriptional patterns of A, B, and D homoeologs

To estimate transcriptional activity and potential tissue specificity of individual PRC1 and PRC2 subunits, we performed transcriptomic analysis using publicly available RNA-sequencing data of 58 bread wheat developmental stages and tissues from the Azhurnaya accession (expVIP database). The transcripts per million (TPM) values were extracted for all of the above-described genes, clustered based on the similarity of their transcriptional profiles over the tissues and visualized in heat maps (Figure 5 and Supplementary Table 2). TPM values were used directly and after log2 transformation, which allows easier analysis of many genes with low transcription levels.

**Figure 5.**
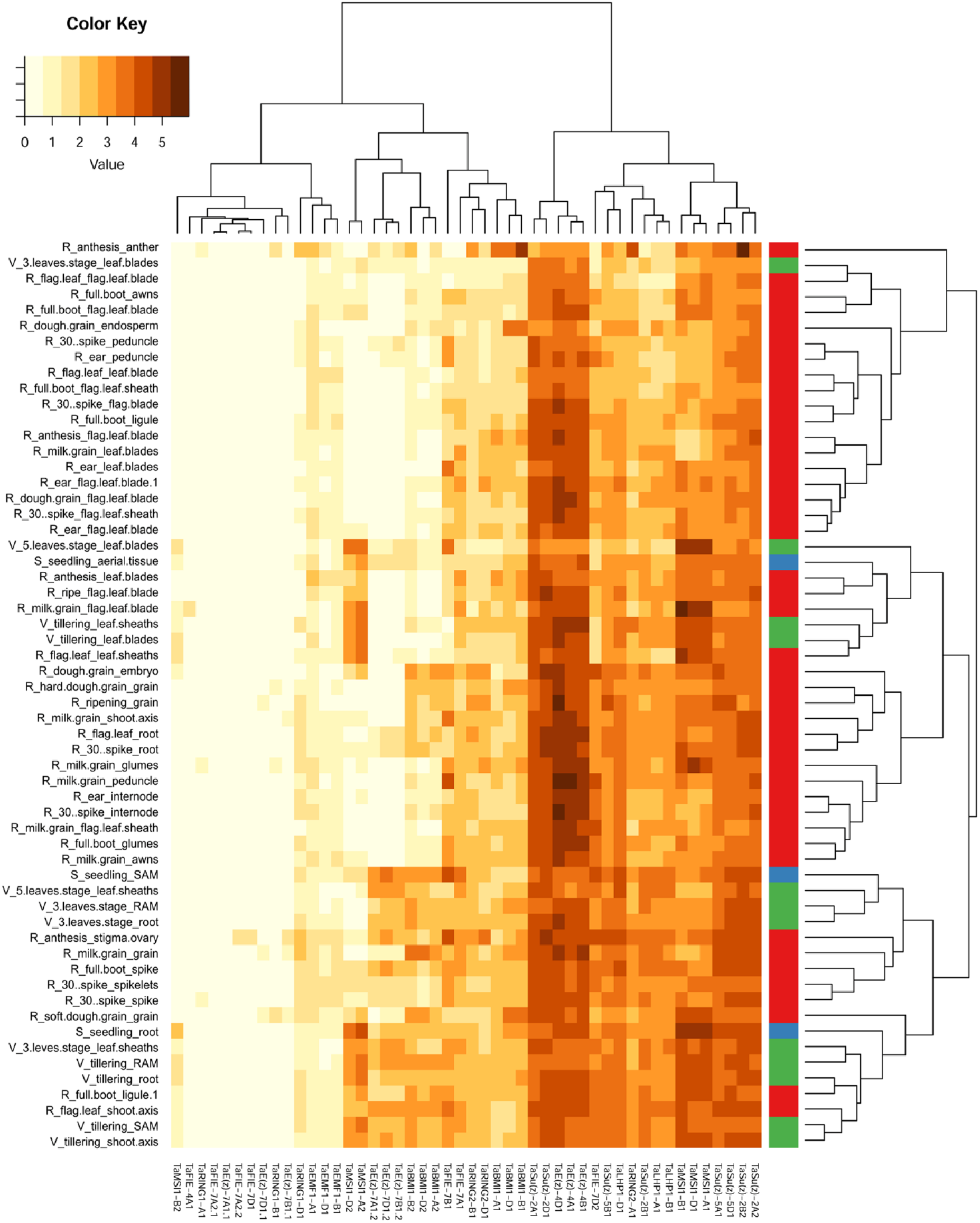
Heat map of PRC1 and PRC2 mRNA levels at different wheat developmental stages. The publicly available RNA-seq data of candidate genes from the cultivar Azhurnaya were clustered based on the transcription profile similarities between the genes (rows) and tissues (columns). Each tissue is characterized as the “high level age_age_tissue”. The high-level stages S – seedling, V – vegetative and R – reproductive are also highlighted by a horizontal color stripe. For a detailed description of the developmental samples and the input values, see Supplementary Table 2. The color key shows transcripts per million (TPM) after log2 transformation.

We found that the homoeologs within the A, B and D subgenomes frequently showed highly similar transcriptional profiles (e.g., *TaE(z)-4A1, B1, D1*; *TaE(z)-7A1.2, B1.2, D1.2*; *TaBMI1-A1, B1, D1*; and *TaBMI1-A2, B2, D2*; *TaMSI1-A1, B1, D1*). This suggests that the developmental regulation established in the progenitor species still exists in the subgenomes of modern wheat and indicates a low degree of functional differentiation between homoeologous gene copies. A possible exception could be *Su(z)-2B2*, which had 61.82 TPM in anthers (R_anthesis_anther), by far the highest value among all genes in the analysis. This mRNA level was 5-fold higher than for its homoeolog *Su(z)-2A2* (TPM 12.39) at the same experimental point. However, both genes showed similar mRNA levels in all other tissues (note that *Su(z)- 2D2* was not found in the *T. aestivum* genome). Although RNA-seq data provided solid support for the transcription of many PRC1 and PRC2 genes, there were also copies that were hardly transcribed in the set of the analyzed tissues, and this held true even for the entire homoeologous group. For example, *TaE(z)-7A1.1, B1.1,* and *D1.1* copies, representing orthologs of *Arabidopsis CLF*, remained practically silent throughout development, while the *TaE(z)* homoeologs on chromosome 4, representing orthologs of *Arabidopsis SWN,* were among the genes with the highest TPM values. A slightly different pattern was observed for *TaMSI1-A2, B2,* and *D2* and *TaMSI1-A1, B1,* and *D1*, representing the tissue-specific and the general MSI groups, respectively. However, such correlations were not universally applicable to all homologs of one PRC1 or PRC2 subunit. Clustering by tissues (log2 plot) revealed three main groups. However, the differences were relatively few. The first two blocks (from left to right in Figure 5) consisted mainly of tissues from plants in the reproductive stage and were characterized by the expression of only specific copies. In contrast, the third cluster contained more tissues from the seedling and vegetative stage plants, which expressed the highest number of PRC1 and PRC2 components.

## DISCUSSION

Plant PcG proteins participate in developmental processes, for example, the transition from the vegetative to generative stage, flowering and seed development (Luo *et al*. 1999; Jiang *et al*. 2008; Xu *et al*. 2018). PcG proteins form groups of Polycomb repressive complexes known as PRC1 and PRC2. PRC2 controls chromatin remodeling through the methylation of histone H3K27 (Margueron and Reinberg 2011). This epigenetic marker of repressed genes is quite common. It has been reported that nearly 4,500 (16 %) genes in *Arabidopsis* carry the repressive mark H3K27me3 (Zhang *et al*. 2007; Farrona *et al*. 2011). In monocots, many genes are also marked with H3K27me3. Interestingly, a significant level of concurrence between the repressive mark H3K27me3 and transcription level was reported in rice, where the majority of the H3K27me3 mark (almost 85 %) is associated with genic regions. In fact, nearly 53 % of H3K27me3-marked genes were expressed, and it was revealed that the gene expression level was correlated with the ratio of H3K4me3/H3K27me3 and H3K27me3/H3K4me3 (He *et al*. 2010). In maize, H3K27me3 is also present mostly in the dense-gene chromosome arms and targets genes with an important regulatory role (Makarevitch *et al*. 2013). Similarly, broad H3K27me3 gene coverage, enhanced in unexpressed genes but unresponsive to the gene expression level, was reported in barley (Baker *et al*. 2014).

The conservation of H3K27me3 targets among plant species was suggested. The targets of H3K27me3 in maize (Makarevitch *et al*. 2013) were compared with genes marked with H3K27me3 in *Arabidopsis* (Lafos *et al*. 2011) and rice (He *et al*. 2010). It was found that 34 % of maize genes that have homologs in *Arabidopsis* were marked with H3K27me3 in both plants. The number of homologous genes marked with H3K27me3 in both monocot species (rice and maize) was almost two times higher than that in *Arabidopsis* (Makarevitch *et al*. 2013). PRC2 also plays a key role in the vernalization response in *Arabidopsis*. Before vernalization, the expression of the major flowering promoter *FLOWERING LOCUS T* (*FT*) is repressed by high levels of *FLC*. Cold treatment triggers PRC2-dependent silencing of *FLC*, which is associated with increased levels of H3K27me3 (Angel *et al*. 2011; Song *et al*. 2012). When *FLC* becomes inactive, the expression of *FT* is initiated and triggers the transition to flowering.

In contrast, temperate cereals were reported to have H3K27me3 marks present at high levels before vernalization (Oliver *et al*. 2009; Xiao *et al*. 2014; Lomax *et al*. 2018), possibly caused by PRC2 activity, as suggested by (Alonso-Peral *et al*. 2011). This may result in chromatin compaction and *VRN1* repression. During the cold period, the H3K27me3 mark disappears, resulting in chromatin remodeling, which may enable the expression of *VRN1*. Consequently, the transition from the vegetative to reproductive stage can occur.

### Identification of PRC2 and PRC1 genes in wheat genomes

The study of molecular mechanisms such as vernalization is hampered by a lack of detailed information about PcG components in bread wheat. Based on homology searches, we identified and located putative PRC2 and PRC1 genes in bread wheat. Most of the subunits were found to be homoeologs in all three wheat subgenomes (A, B and D).

Chromosomal positions of the wheat PRC2 components corresponded with the previously reported PRC2 genes in barley (Kapazoglou *et al*. 2010). Interestingly, several paralogs were located within the same chromosome, and paralogs located on different chromosomes were also found. These multiple locations could be explained by the allohexaploid nature of the wheat genome, which has undergone frequent chromosomal rearrangements. The comparison between individual paralogs also revealed shortened proteins (Supplementary Figure S1) with low expression levels. These findings indicate the possible alteration and/or subfunctionalization of the genes. We also found paralogs that differ at the distance between individual copies. *TaSu(z)-2A1* and *TaSu(z)-2A2* are separated by more than 1.1 Mb, while two copies of *TaFIE* genes (*TaFIE-7A2.1* and *TaFIE-7A2.2*) are separated only by a region of 37 kb (Supplementary Table 1), which indicates that different mechanisms contribute to gene duplications in wheat. Unfortunately, their expression level based on the expVIP database is minimal.

Interestingly, E(z) paralogs were identified on chromosome groups 4 and 7. A translocation between chromosomes 4 and 7 has been reported (Hernandez *et al*. 2012; The International Wheat Genome Sequencing Consortium (IWGSC) *et al*. 2018). Briefly, the structure of present-day wheat chromosome 4 is an illustrative example of dynamic chromosomal rearrangements within the allohexaploid wheat genome. The final composition of the chromosome resulted from pericentric inversion of the ancient long arm, which became a modern short arm, and the subsequent translocation from 5AL and 7BS completed the rearrangement of the chromosome. In agreement with this, the copy of the *TaFIE-4A1* gene maintained a closer phylogenetic relationship to the homologs on the 7AS chromosome (*TaFIE-7A2.1* and *TaFIE-7A2.2*) (Figure 3A).

Moreover, the phylogenetic analysis revealed that genes on chromosome 4 are putative orthologs to *AtSWN*, while genes on chromosome 7 are closer to *AtCLF*. Protein alignment of conserved domains from *Arabidopsis* SWN and CLF with domains from TaE(z) revealed nine amino acid changes in the catalytic SET domain correlating with the differentiation of SWN- and CLF-like TaE(z) proteins, which further supports this hypothesis (Supplementary Figure S2). CLF and SWN are largely functionally redundant in *Arabidopsis,* and their simultaneous knockout in plants results in the production of callus-like structures containing somatic embryos (Mozgová *et al*. 2017). Currently, the extent of functional redundancy between the *TaSWN-like* and *TaCLF-like* groups is unknown, but *TaSWN-like* homoeologs are more strongly expressed copies than *TaCLF-like* homoeologs, which contrasts the pattern in *Arabidopsis* (Schmid *et al*. 2005). There was also a substantial difference in mRNA levels (up to 11-fold) between *CLF-like* paralogs on chromosome 7. This could indicate that the cis-regulatory elements of some copies were either mutated or lost. Future experiments will show whether such copies may be either subfunctionalized at the tissue-specific level or progressing towards removal from the bread wheat genome. The analysis of the expression profile showed that not all paralogs representing individual core components were expressed similarly, but there was always at least one gene with a high expression level. This could be because the paralog sequences were not identical (Supplementary Figure S1); therefore, their function and expression could be altered.

Unlike the identification of LHP1, RING1 and BMI1, which assemble the core components of plant PRC1, the identification of other plant-specific proteins that may be part of this complex was more difficult. The chemical properties and functions of EMF1 are similar to those of Psc in *Drosophila* and its ortholog, BMI1, in *Arabidopsis* (Beh *et al*. 2012). The poorly conserved sequence of EMF1 does not display significant homology with any other proteins of known function (Aubert *et al*. 2001). There are no annotated domains in EMF1, but five conserved motifs shared by the entire EMF1 orthologous group were predicted (Calonje *et al*. 2008; Berke and Snel 2015). Despite the fact that the presence of EMF1 was shown in both monocots and eudicots (Aubert *et al*. 2001; Calonje *et al*. 2008; Berke and Snel 2015), no direct homolog was found in *T. aestivum* using the EMF1 protein sequence from *Arabidopsis* for homology searches. Therefore, we had to use a sequence of a monocot plant (maize), which could imply that EMF1 is less conserved among dicots and monocots. *AtVRN1*, which was assigned in previous studies to PRC1 (Mylne *et al*. 2006; Holec and Berger 2011), was shown to be absent in monocots (Berke and Snel 2015). In *Arabidopsis*, *AtVRN1* plays an important role in vernalization. It should be emphasized that the *VERNALIZATION1* (*VRN1*) gene in wheat is not related to *VRN1* in *Arabidopsis* but is homologous to *APETALA1, CAULIFLOWER* and *FRUITFUL* (*AP1, CAL* and *FUL*) with no role in *Arabidopsis* vernalization (Yan *et al*. 2003). However, when the AtVRN1 protein sequence from *Arabidopsis* was used for a homology search in wheat, similar proteins, whose genes are located on chromosomes 5A, 5B and 5D, were obtained. These proteins contain four B3 domains, whereas the AtVRN1 protein in *Arabidopsis* contains only two domains. In summary, all core subunits of PRC1 (consisting of LHP1, RING1 and BMI1) in monocots were identified in bread wheat. The identification of the plant-specific proteins EMF1 and VRN1 remains less evident. Individual subunits of PRC1 also share conserved protein domains between paralogs, but not all paralogs had the same expression level, indicating differentiation at the cis-regulatory level.

In conclusion, the identification of individual PcG components in bread wheat will help to reveal the molecular mechanisms of important biological processes. More detailed studies (expression studies, sequence variation among wheat cultivars, etc.) will be necessary to reveal the possible functional divergence of single genes, including paralogs, and their putative role in the formation of Polycomb repressive complexes affecting plant development.

## Author contributions

ZM conceived the study, BS carried out bioinformatics analysis and technical preparation of the manuscript. RC reconstructed nucleotide sequences from scaffolds and performed phylogenetic analysis. AP analyzed the RNA-seq data. JS contributed to the interpretation of the results. All authors discussed the results and wrote the manuscript.

## Supplementary material

*Supplementary Table 1*

List of bread wheat accessions for individual PcG components and comparision of their chromosome locations with T. urartu, T. dicoccoides and H. vulgare homologs

*Supplementary Table 2*

RNA-seq data of bread wheat PcG genes used for transcriptomic analysis

*Supplementary Table 3*

Protein sequences of PcG components used for phylogenetic analysis

*Supplementary Figure 1*

Protein alignments of plant PRC1 and PRC2 core components

*Supplementary Figure 2*

Protein alignment of E(z) homologs showing nine aminoacid exchanges in SET domain corresponding with division of TaE(z) paralogs into SWN- and CLF-like groups

## ACKNOWLEDGMENTS

We would like to thank Dr. Martin Mascher (IPK, Gatersleben, Germany) for sharing unpublished sequencing data of barley genome.

