## Supplementary Figure 1 for "Identification of Polycomb Repressive Complex 1 and 2 Core Components in Hexaploid Bread Wheat"

**Supplementary Figure 1.** Protein alignments of plant PRC1 core components LHP1 **(A),** BMI1 **(B)** and RING1 **(C)** and PRC2 core components E(z) **(D)**, FIE **(E)**, MSI **(F)** and Su(z) **(G)** from *Arabidopsis thaliana* (At), *Hordeum vulgare* (HORVU), *Triticum dicoccoides* (TRIDC) and *Triticum aestivum* (Ta). Conserved protein domains are highlighted. CD; chromo domain, CSD; chromo shadow domain.

**A**

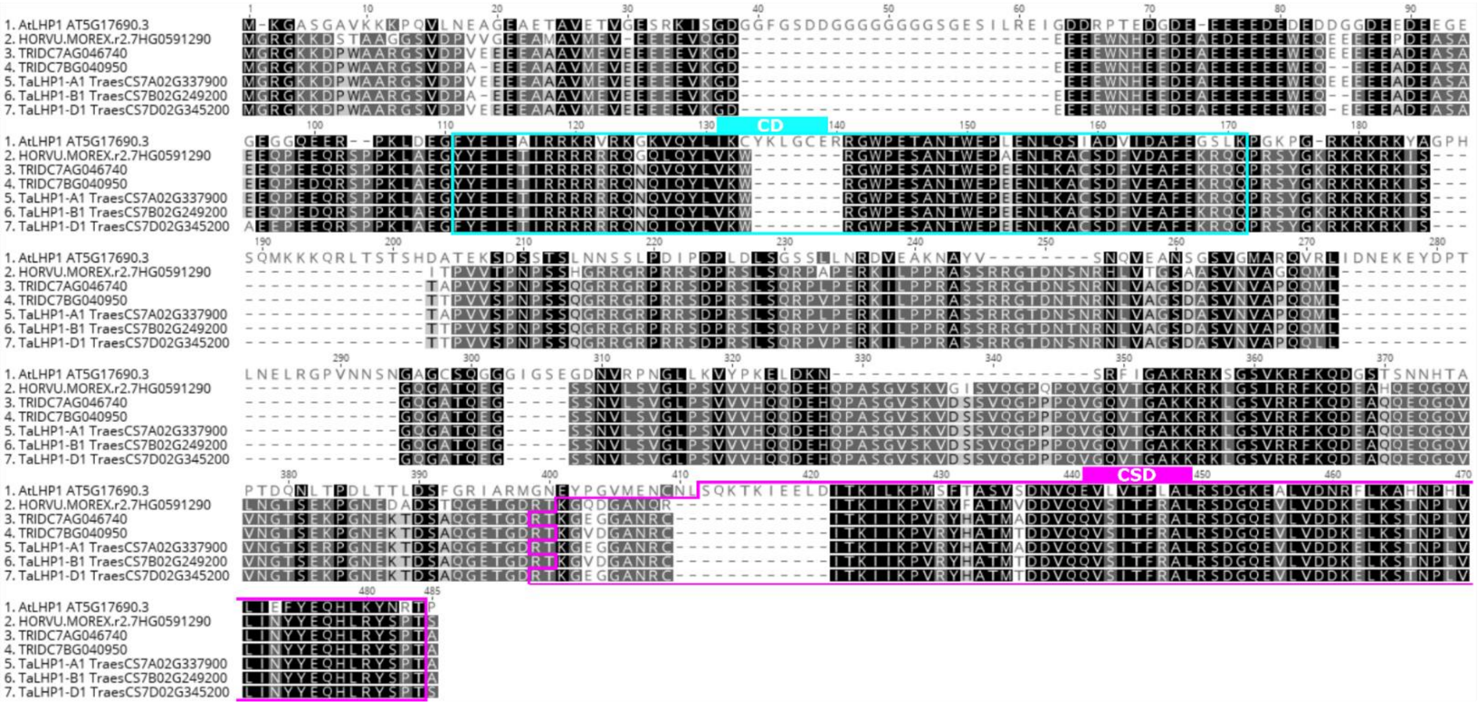

**B**

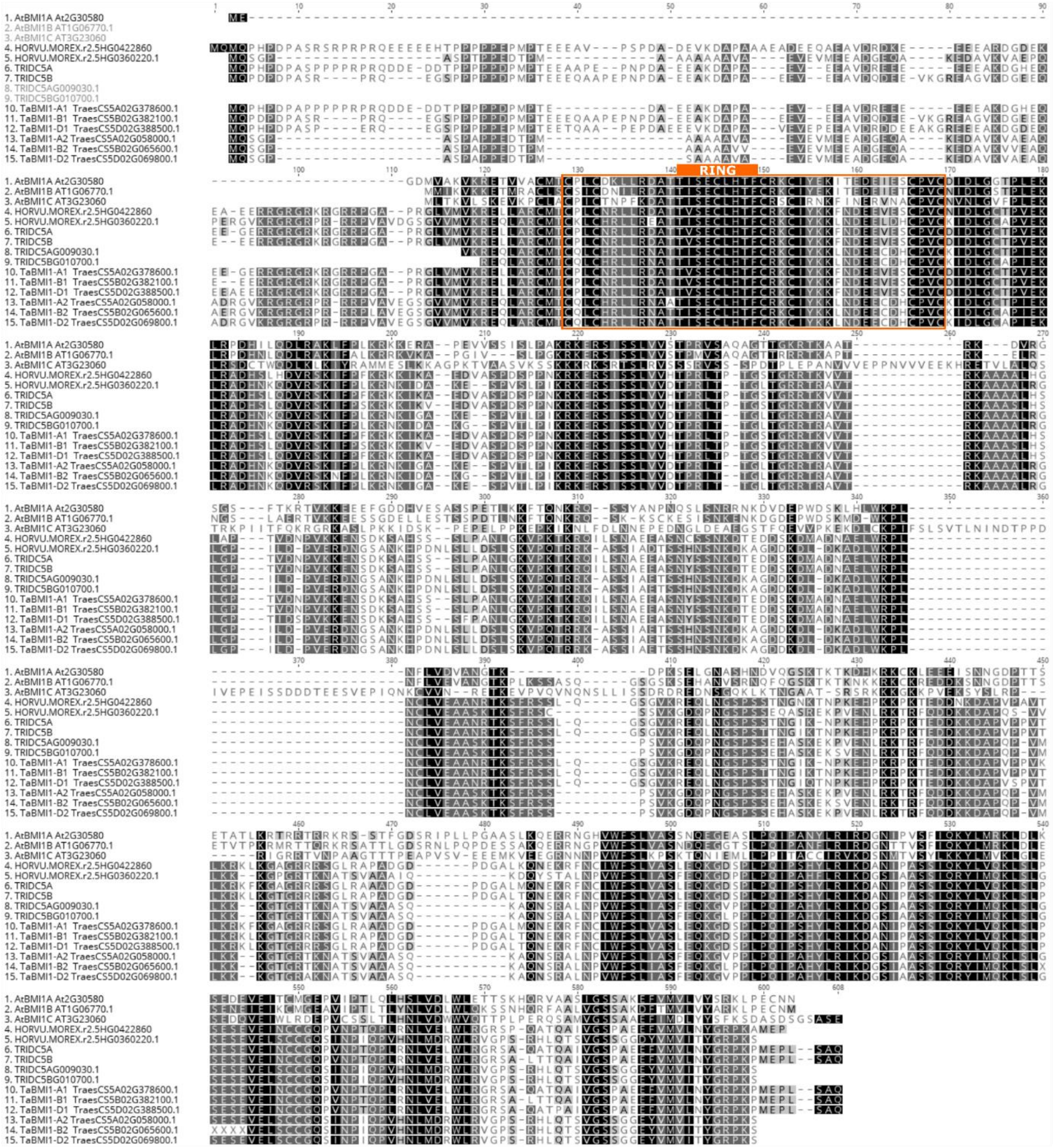
RING
1. AtBMI1A At2G30580
2. AtBMI1B AT1G06770.1
3. AtBMI1C AT3G23060
4. HORVU.MOREX.r2.5HG0422860
5. HORVU.MOREX.r2.5HG0360220.1
6. TRIDC5A
7. TRIDC5B
8. TRIDC5AG009030.1
9. TRIDC5BG010700.1
10. TaBMI1-A1 TraesCS5A02G378600.1
11. TaBMI1-B1 TraesCS5B02G382100.1
12. TaBMI1-D1 TraesCS5D02G388500.1
13. TaBMI1-A2 TraesCS5A02G058000.1
14. TaBMI1-B2 TraesCS5B02G065600.1
15. TaBMI1-D2 TraesCS5D02G069800.1

C

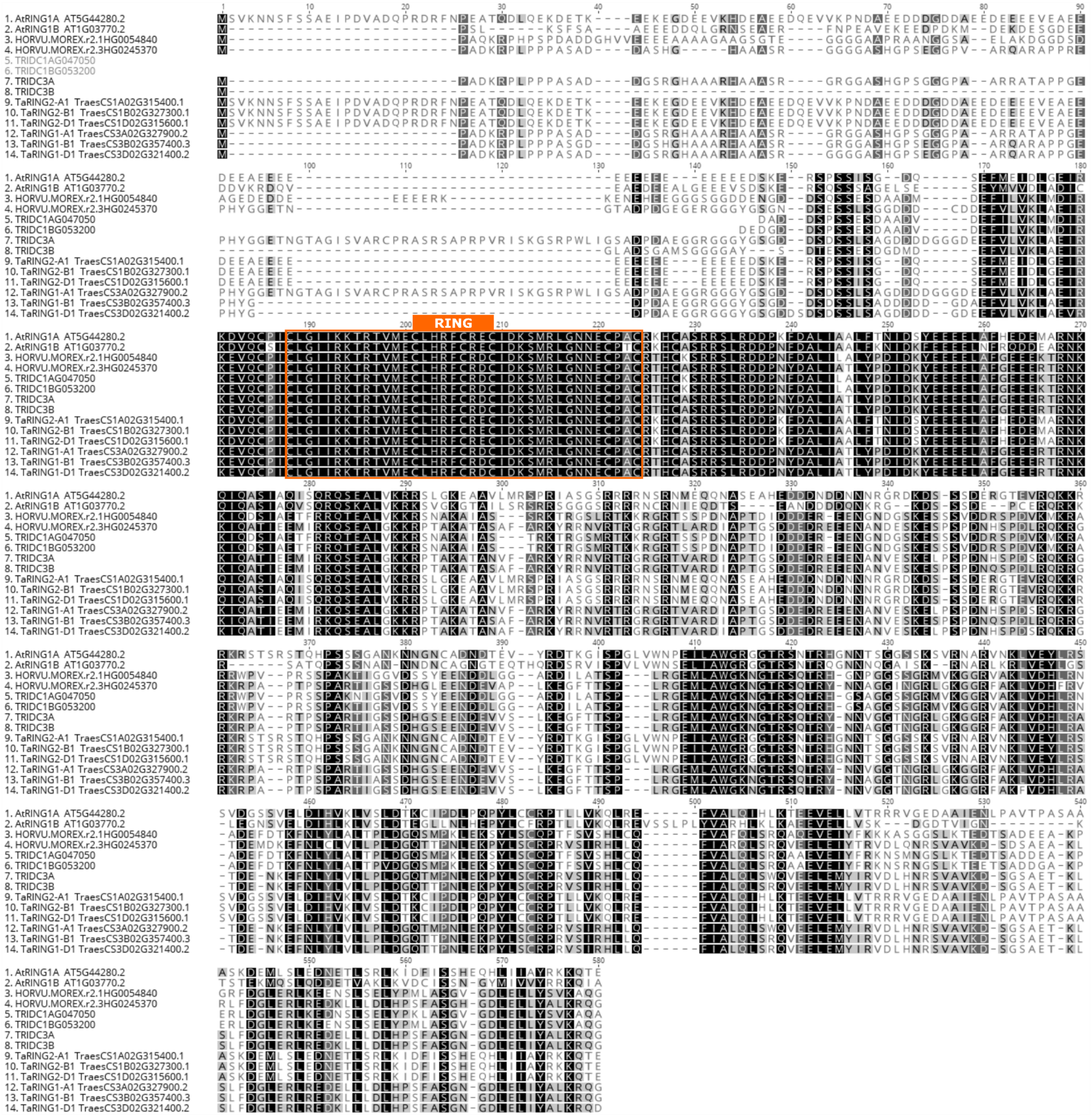
1. AtRING1A AT5G44280.2
2. AtRING1B AT1G03770.2
3. HORVU.MOREX.r2.1HG0054840
4. HORVU.MOREX.r2.3HG0245370
5. TRIDC1AG047050
6. TRIDC1BG053200
7. TRIDC3A
8. TRIDC3B
9. TaRING2-A1 TraesCS1A02G315400.1
10. TaRING2-B1 TraesCS1B02G327300.1
11. TaRING2-D1 TraesCS1D02G315600.1
12. TaRING1-A1 TraesCS3A02G327900.2
13. TaRING1-B1 TraesCS3B02G357400.3
14. TaRING1-D1 TraesCS3D02G321400.2
RING

### D (part1)

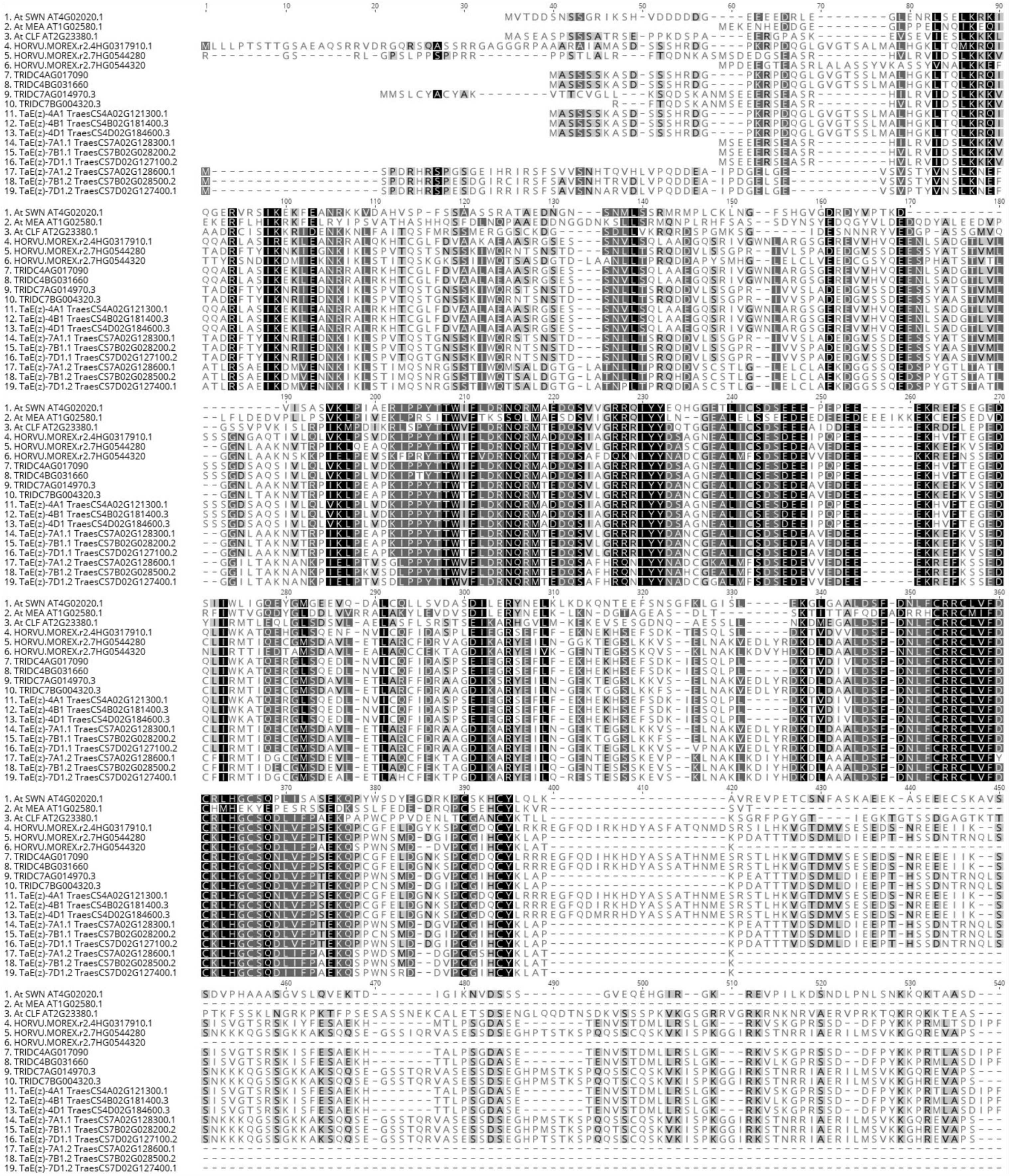
1. At SWN AT4G02020.1
2. At MEA AT1G02580.1
3. At CLF AT2G23380.1
4. HORVU.MOREX.r2.4HG0317910.1
5. HORVU.MOREX.r2.7HG0544280
6. HORVU.MOREX.r2.7HG0544320
7. TRIDC4AG017090
8. TRIDC4BG031660
9. TRIDC7AG014970.3
10. TRIDC7BG004320.3
11. TaE(z)-4A1 TraesCS4A02G121300.1
12. TaE(z)-4B1 TraesCS4B02G181400.3
13. TaE(z)-4D1 TraesCS4D02G184600.3
14. TaE(z)-7A1.1 TraesCS7A02G128300.1
15. TaE(z)-7B1.1 TraesCS7B02G028200.2
16. TaE(z)-7D1.1 TraesCS7D02G127100.2
17. TaE(z)-7A1.2 TraesCS7A02G128600.1
18. TaE(z)-7B1.2 TraesCS7B02G028500.2
19. TaE(z)-7D1.2 TraesCS7D02G127400.1

**D (part2)**

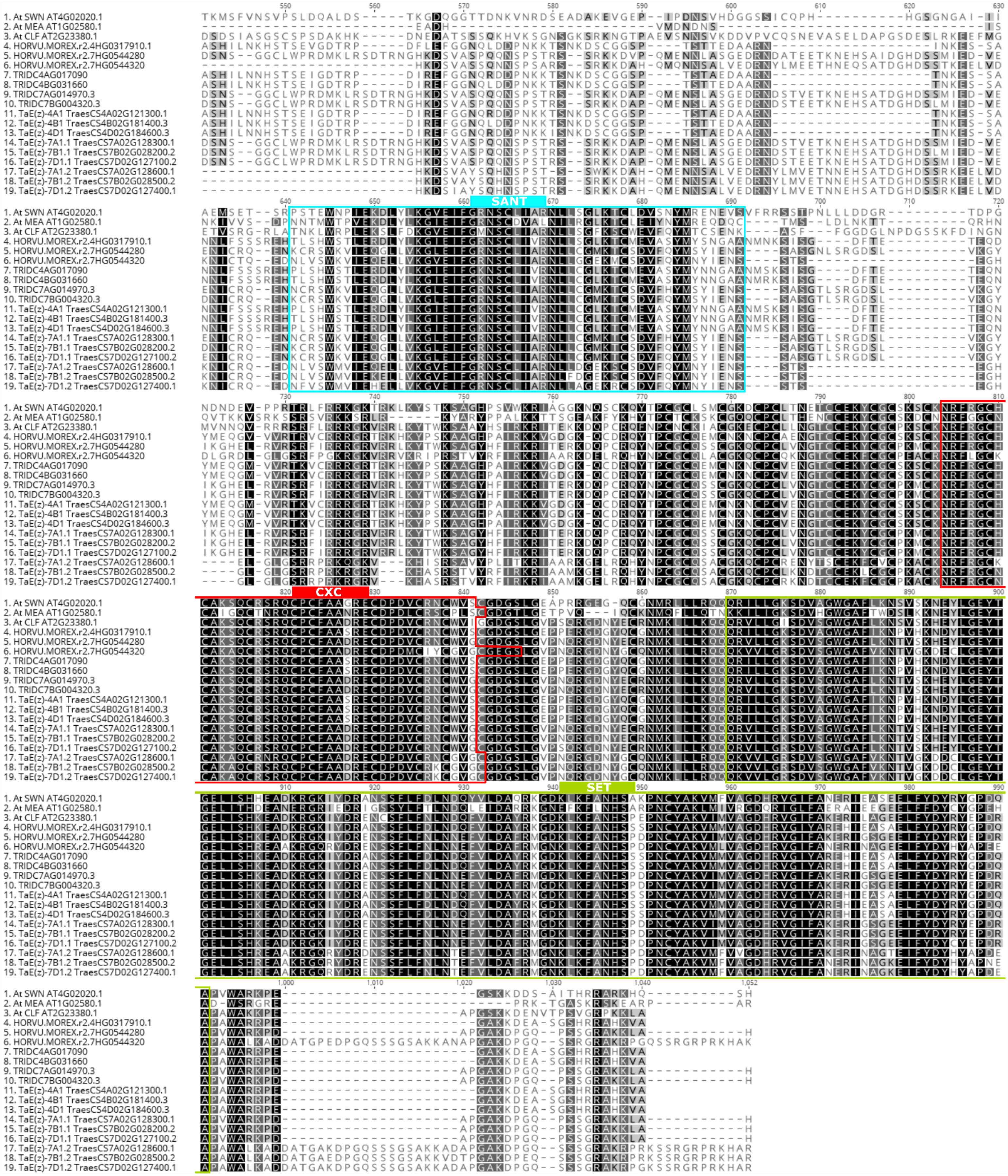
1. At SWN AT4G02020.1
2. At MEA AT1G02580.1
3. At CLF AT2G23380.1
4. HORVU.MOREX.r2.4HG0317910.1
5. HORVU.MOREX.r2.7HG0544280
6. HORVU.MOREX.r2.7HG0544320
7. TRIDC4AG017090
8. TRIDC4BG031660
9. TRIDC7AG014970.3
10. TRIDC7BG004320.3
11. TaE(z)-4A1 TraesCS4A02G121300.1
12. TaE(z)-4B1 TraesCS4B02G181400.3
13. TaE(z)-4D1 TraesCS4D02G184600.3
14. TaE(z)-7A1.1 TraesCS7A02G128300.1
15. TaE(z)-7B1.1 TraesCS7B02G028200.2
16. TaE(z)-7D1.1 TraesCS7D02G127100.2
17. TaE(z)-7A1.2 TraesCS7A02G128600.1
18. TaE(z)-7B1.2 TraesCS7B02G028500.2
19. TaE(z)-7D1.2 TraesCS7D02G127400.1
SANT
CXC
SET

E

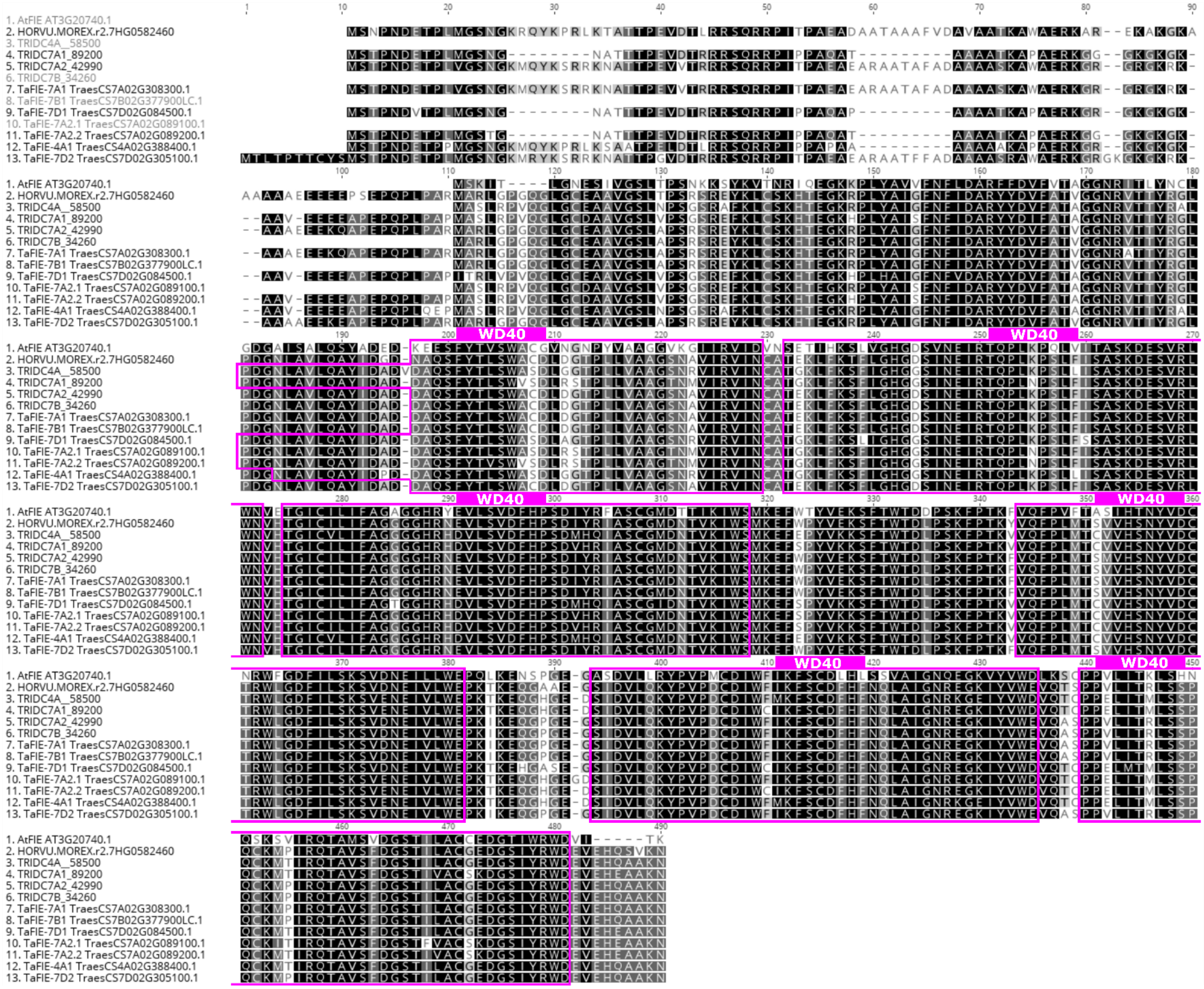
1. AtFIE AT3G20740.1
2. HORVU.MOREX.r2.7HG0582460
3. TRIDC4A_58500
4. TRIDC7A1_89200
5. TRIDC7A2_42990
6. TRIDC7B_34260
7. TaFIE-7A1 TraesCS7A02G308300.1
8. TaFIE-7B1 TraesCS7B02G377900LC.1
9. TaFIE-7D1 TraesCS7D02G084500.1
10. TaFIE-7A2.1 TraesCS7A02G089100.1
11. TaFIE-7A2.2 TraesCS7A02G089200.1
12. TaFIE-4A1 TraesCS4A02G388400.1
13. TaFIE-7D2 TraesCS7D02G305100.1
WD40
WD40
WD40
WD40
WD40
WD40

**F**

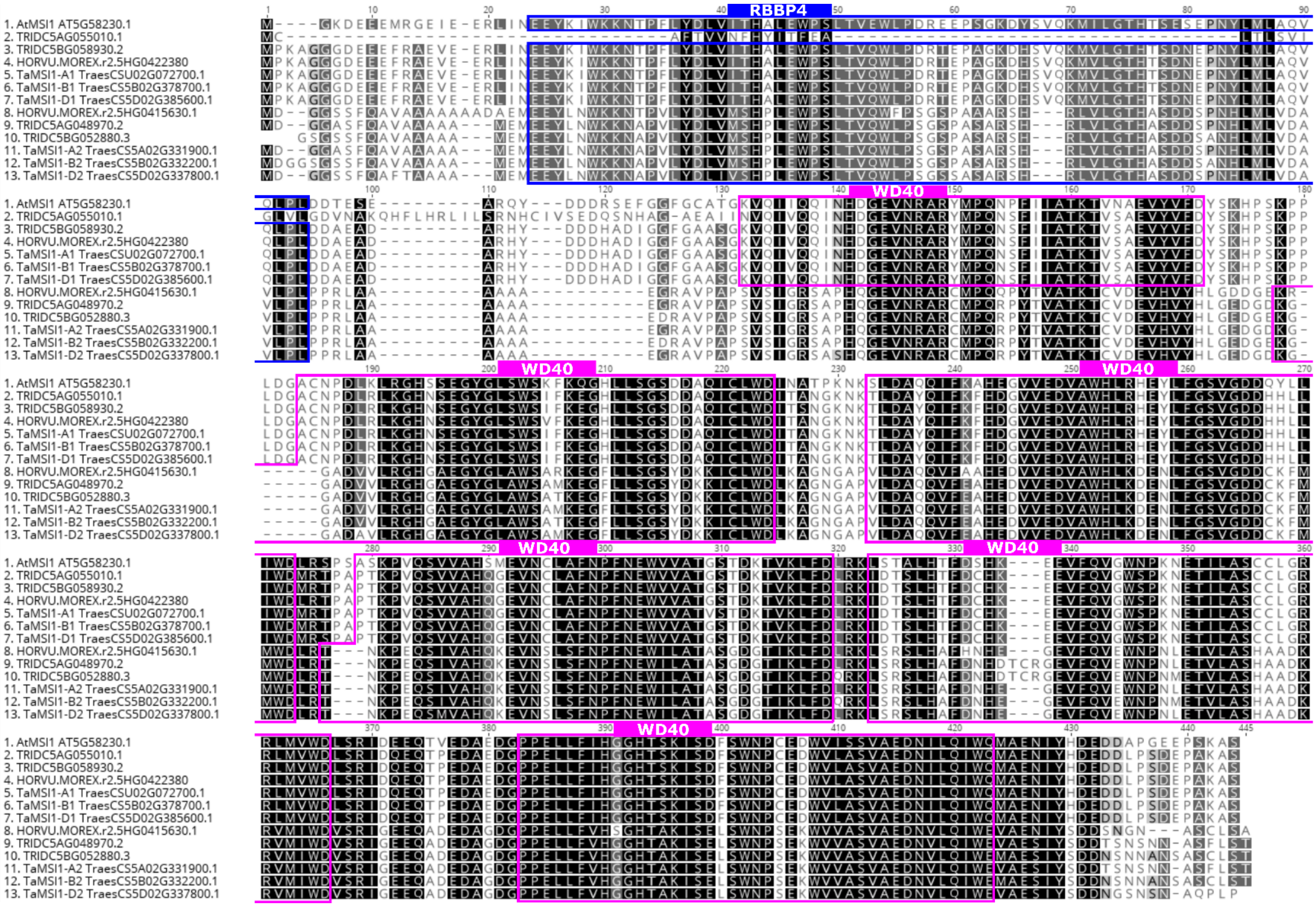
RBBP4
WD40
WD40
WD40
WD40
WD40
WD40
1. AtMSI1 AT5G58230.1
2. TRIDC5AG055010.1
3. TRIDC5BG058930.2
4. HORVU.MOREX.r2.5HG0422380
5. TaMSI1-A1 TraesCSU02G072700.1
6. TaMSI1-B1 TraesCS5B02G378700.1
7. TaMSI1-D1 TraesCS5D02G385600.1
8. HORVU.MOREX.r2.5HG0415630.1
9. TRIDC5AG048970.2
10. TRIDC5BG052880.3
11. TaMSI1-A2 TraesCS5A02G331900.1
12. TaMSI1-B2 TraesCS5B02G332200.1
13. TaMSI1-D2 TraesCS5D02G337800.1

**G (part1)**

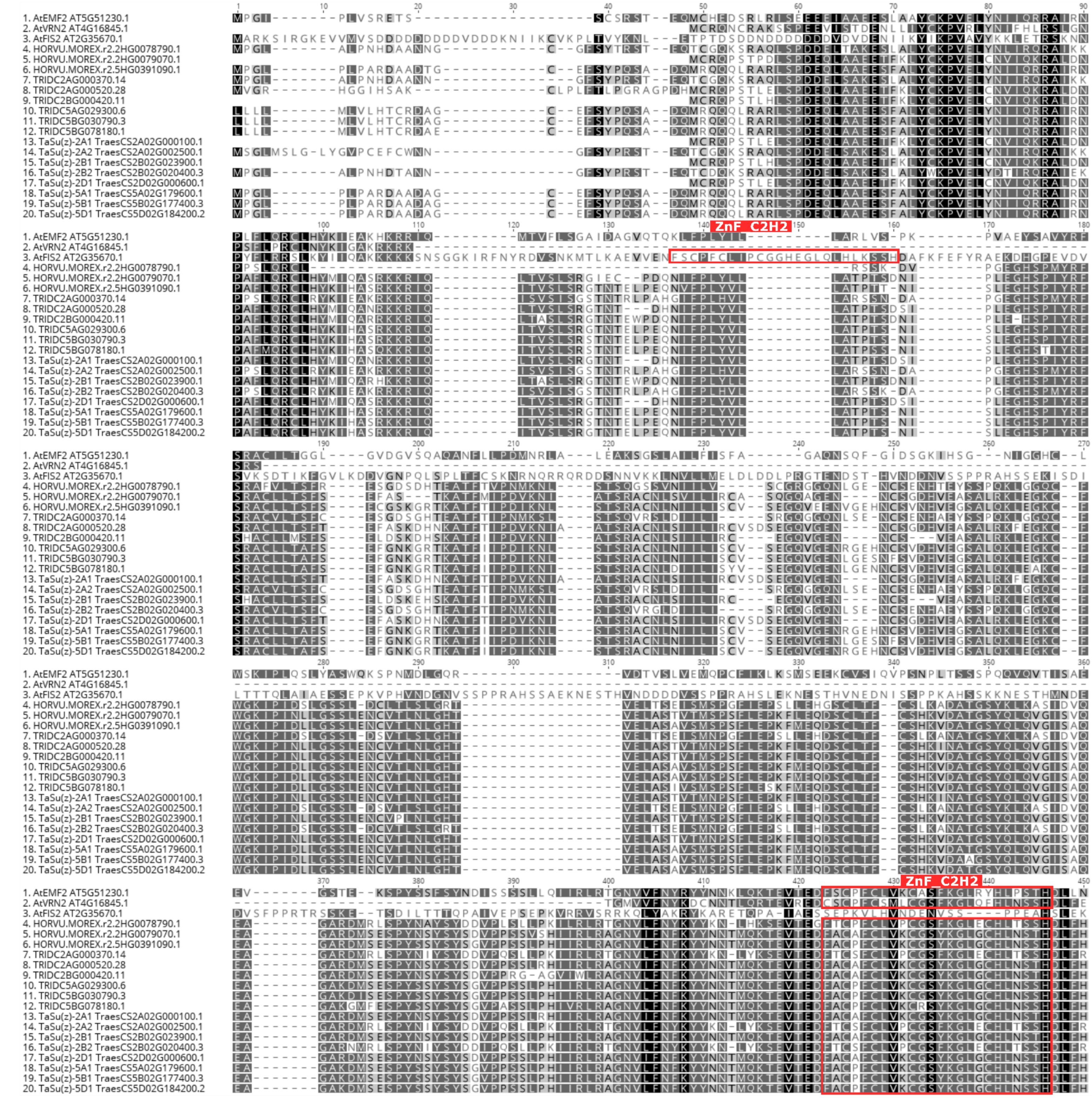
ZnF_C2H2
ZnF_C2H2
1. AtEMF2 AT5G51230.1
2. AtVRN2 AT4G16845.1
3. AtFIS2 AT2G35670.1
4. HORVU.MOREX.r2.2HG0078790.1
5. HORVU.MOREX.r2.2HG0079070.1
6. HORVU.MOREX.r2.5HG0391090.1
7. TRIDC2AG000370.14
8. TRIDC2AG000520.28
9. TRIDC2BG000420.11
10. TRIDC5AG029300.6
11. TRIDC5BG030790.3
12. TRIDC5BG078180.1
13. TaSu(z)-2A1 TraesCS2A02G000100.1
14. TaSu(z)-2A2 TraesCS2A02G002500.1
15. TaSu(z)-2B1 TraesCS2B02G023900.1
16. TaSu(z)-2B2 TraesCS2B02G020400.3
17. TaSu(z)-2D1 TraesCS2D02G000600.1
18. TaSu(z)-5A1 TraesCS5A02G179600.1
19. TaSu(z)-5B1 TraesCS5B02G177400.3
20. TaSu(z)-5D1 TraesCS5D02G184200.2

**G (part2)**

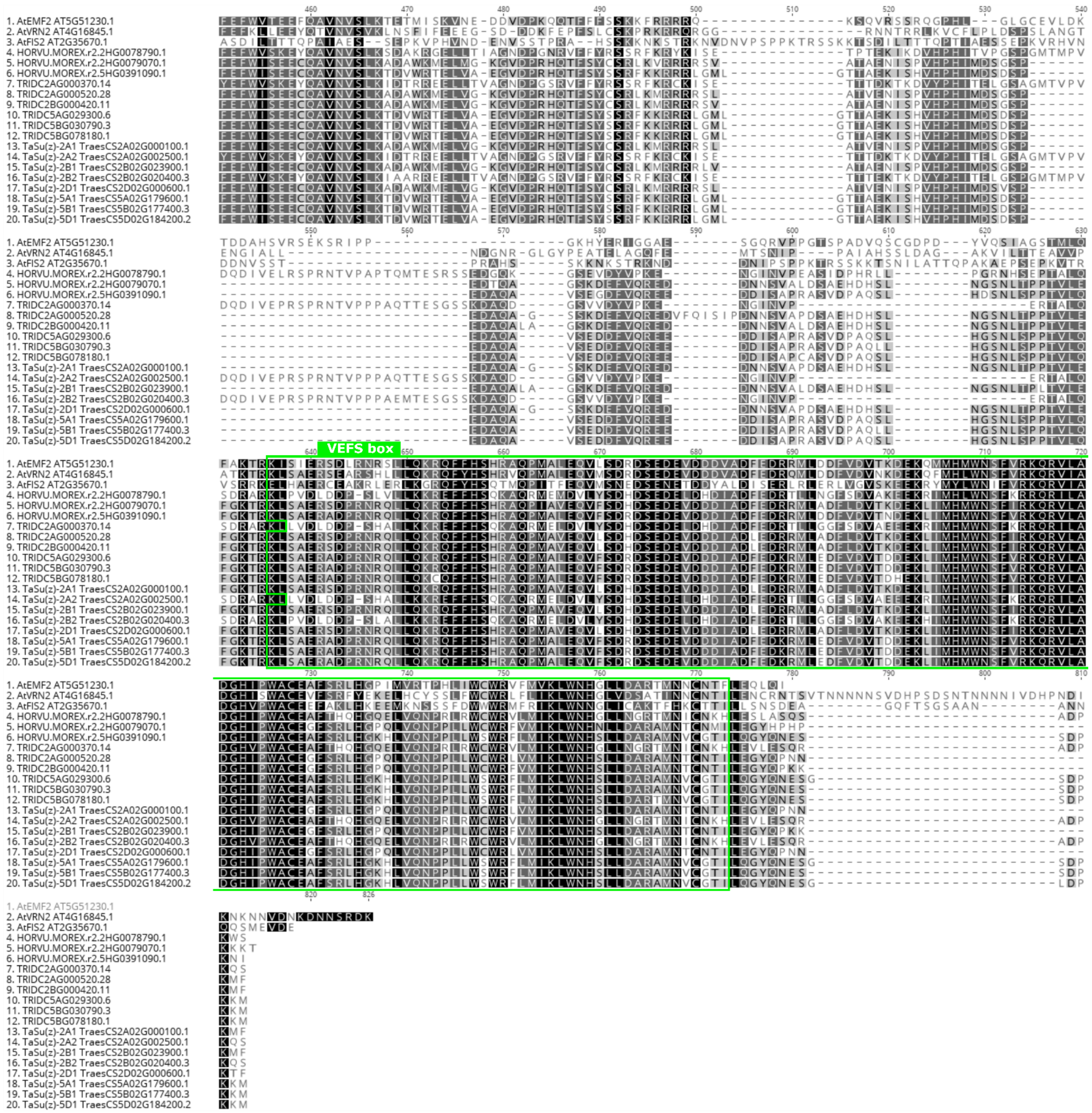
VEFS box
1. AtEMF2 AT5G51230.1
2. AtVRN2 AT4G16845.1
3. AtFIS2 AT2G35670.1
4. HORVU.MOREX.r2.2HG0078790.1
5. HORVU.MOREX.r2.2HG0079070.1
6. HORVU.MOREX.r2.5HG0391090.1
7. TRIDC2AG000370.14
8. TRIDC2AG000520.28
9. TRIDC2BG000420.11
10. TRIDC5AG029300.6
11. TRIDC5BG030790.3
12. TRIDC5BG078180.1
13. TaSu(z)-2A1 TraesCS2A02G000100.1
14. TaSu(z)-2A2 TraesCS2A02G002500.1
15. TaSu(z)-2B1 TraesCS2B02G023900.1
16. TaSu(z)-2B2 TraesCS2B02G020400.3
17. TaSu(z)-2D1 TraesCS2D02G000600.1
18. TaSu(z)-5A1 TraesCS5A02G179600.1
19. TaSu(z)-5B1 TraesCS5B02G177400.3
20. TaSu(z)-5D1 TraesCS5D02G184200.2
