## Supplementary Figure 2 for "Identification of Polycomb Repressive Complex 1 and 2 Core Components in Hexaploid Bread Wheat"

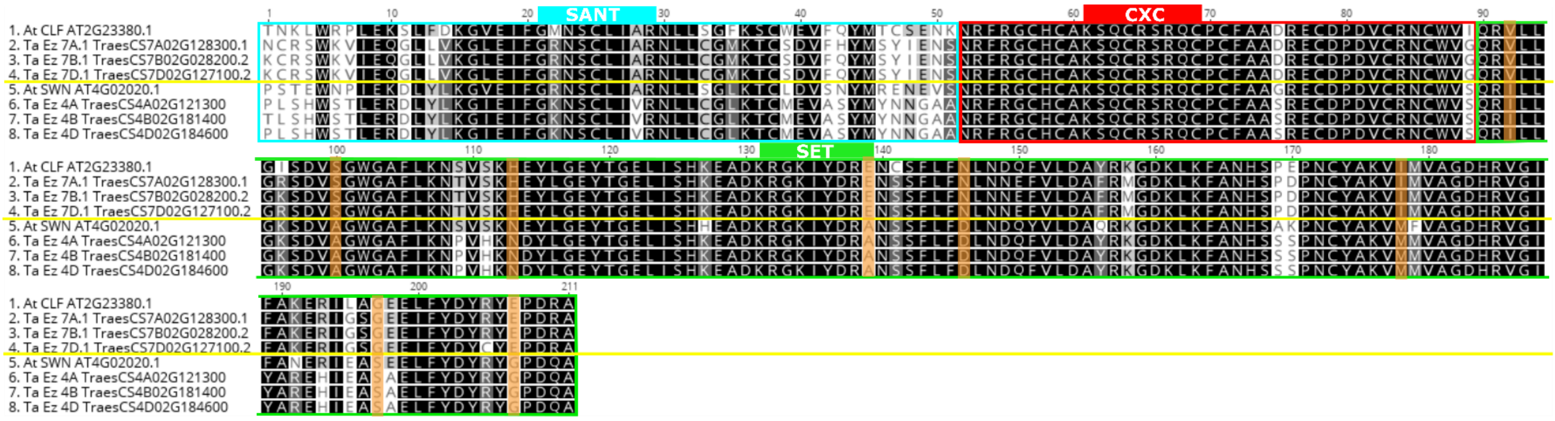


**Supplementary Figure 2.** Nine aminoacid exchanges in SET domain corresponding with division of TaE(z) paralogs into SWN- and CLF-like groups are highlighted with orange color.
